# Synthetic extremophiles: Species-specific formulations for microbial therapeutics and beyond

**DOI:** 10.1101/2022.11.30.518573

**Authors:** Miguel Jimenez, Johanna L’Heureux, Emily Kolaya, Kyle B. Martin, Zachary Villaverde, Afeefah Khazi-Syed, Qinhao Cao, Benjamin Muller, James D. Byrne, Giovanni Traverso

## Abstract

Microorganisms have been used for millennia to produce food and medicine and are now being developed as products themselves to treat disease and boost crop production. However, as required for these new applications, maintaining high viability throughout manufacturing, transportation and use remains a significant challenge requiring sophisticated cold-chains and packaging. In fact, we found that commercial microbial products (probiotics) provide a poor solution to this challenge, in particular for key industrial organisms like *E. coli*. To overcome this technological gap, we report the development of synthetic extremophiles of industrially important gram-negative bacteria (*E. coli* Nissle 1917, *Ensifer meliloti*), gram positive bacteria (Lactobacillus *plantarum*) and yeast (*Saccharomyces boulardii*). Specifically, we developed a high throughput pipeline to define species-specific materials that allow these organisms to survive drying, elevated temperatures, organic solvents and even ionizing radiation. We enhanced the stability of *E*.*coli* Nissle 1917 by >4 orders of magnitude over commercial formulations and demonstrate the capacity to remain viable while undergoing tableting and pharmaceutical methodologies involving organic solvents. The development of synthetic materials-based enhanced stabilization stands to transform our capacity to apply micro-organisms in extreme environments including those found on Earth as well as in space.

**One-Sentence Summary:** Fragile therapeutic bacteria can be made to survive the manufacturing extremes normally reserved for small molecule drugs.

## Main Text

Microorganisms have been central to human technological progress and continue to be key in wide ranging fields from food production (e.g. baked goods) to biologics manufacturing (e.g. synthetic insulin)^1^. However, by and large, microbial cells are kept alive only during the manufacturing process and are destroyed, deactivated, or removed from the final product.

However, through the advent of culture-independent sequencing techniques and synthetic biology, the pharmaceutical, agricultural and space health fields have now turned to developing live microorganisms as the final product to cure disease^2^, enhance crop production^3^ and for on-demand bioproduction^4^.

Critical to these new microbial technologies is maintenance of high cell viability throughout the entire life cycle of the product. To achieve this, researchers and commercial companies have turned to well-established microbial cryopreservation methods^5^ or to organisms that have natural extremophilic properties (e.g. dehydration-resistance, acid-resistance)^6^. However, the need for select organisms or costly, inflexible cold chains severely limits the use case of live microbial products.

An ideal solution would be dry, shelf-stable microbial materials that are simple to package, ship, and use. While there has been extensive research on the lyopreservation of microorganisms, these previous studies have primarily focused on storage of microorganisms in culture collections which only require “acceptable” levels of viability (i.e., just enough viable cells for recovery through culture amplification)^7,8^. Furthermore, while previous studies have defined sugars and peptides with high glass transition temperatures as generally good stabilizers, these studies have also shown that specific viability results vary widely depending on the microorganism in question^7,8^.

Therefore, in this work, we first evaluated available commercial examples of dried microbial products (i.e., probiotics) to understand the current state of microbial stabilization. We found generally low viabilities with particularly poor results for gram-negative bacteria (e.g., *E. coli*). To fill this gap, we developed material-based synthetic extremophiles that are shelf-stable without refrigeration. We further show that that these stabilized microorganisms can withstand the extreme conditions encountered in pharmaceutical manufacturing pipelines (i.e., heat, pressure, solvents). Finally, we demonstrate that this stabilization approach maintains the functional capacity of the microorganisms in a bioluminescence and plant root nodulation assay.

### Survey of commercial probiotic products

A salient example of dried microbial products are the current commercially available probiotics marketed for human use that line pharmacy shelves. These products represent the current state of the art of microbial stabilization for commercial applications. However, when we surveyed the viable cell counts (colony forming units, CFUs) across a range of off-the-shelf probiotics (Table S1), we found only 7 in 13 products contained viable cell counts at or higher than the promised amount on the label (Fig. 1, S1), with a mean (geo.) viability of just ∼21% of that promised. Nevertheless, when we assessed the total cell count microscopically (dead + alive) all products contained cell totals above those promised (Fig. S1A), suggesting a loss of viability during and following manufacture. When we compared the viable cell count to this microscopically determined total cell count, only 1 in 13 products had a viability greater than 40%, with 6 in 13 products with viabilities lower than 2% and an overall mean viability of just 1.9% of total cells (Fig. 1).

**Fig. 1.**
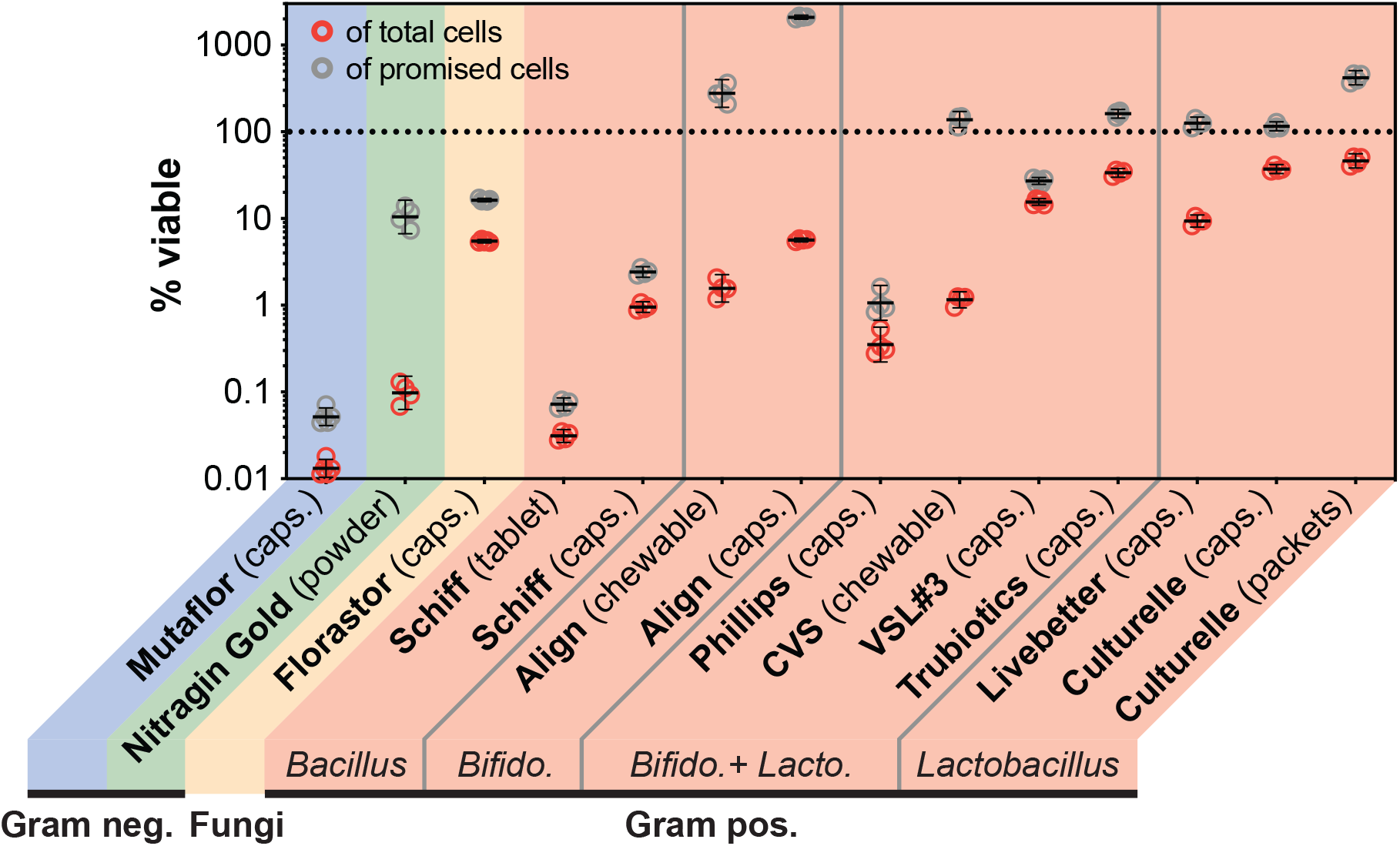
Survey of commercially available dry formulations of microbial probiotics. We quantified the viability of a wide representative range of commercial probiotic products (Table S1). Viable cell counts were defined as the number of colony forming units (CFUs) assessed through dilution plating on the appropriate solid medium and quantified via programmatic image analysis (see Methods). Percent viable rates relative to promised cells were computed by dividing the CFU count by the CFU counts printed on the product label. Percent viable rates relative to total cells were computed by dividing the CFU counts by the number of total countable cells determined via automated fluorescence cytometry. N = 4 or 5 independent dosage forms. Geometric means and geometric 95% confidence intervals plotted in black over individual replicates. Broad phylogenetic classes of the component microorganisms are noted in color and specific genera noted in italics (see Fig. S1 for detailed make up). neg. = negative; pos. = positive; Bifido. = Bifidobacterium; Lacto. = Lactobacillus.

Beyond this overall low viability rate, we also found a lack of variety in terms of microbial species identity (Fig. S1B). A majority (9 of 13) of products used microorganisms from just two groups (*Lactobacillus, Bifidobacterium*) which encompassed all the products with higher viabilities (Fig. S1B). There was only one commercially available representative in the large clade of gram-negative bacteria (Mutaflor, *Escherichia coli* Nissle 1917) and it had the lowest viability (0.05% of promised and 0.01% of total cells). To corroborate this finding, we found one additional commercial gram-negative product marketed for plant-use (Nitragin Gold, *Ensifer meliloti*) and found it also had a low viability (10% of promised and 1% of total cells) (Fig. 1).

Furthermore, to evaluate the intrinsic extremophilic qualities of these products, we stress tested them by exposing them to 50 °C for 24 hours. We found nearly all the products retained an acceptable viability of >10% relative to non-stressed samples (Fig. S2). However, the *E. coli* Nissle 1917 product Mutaflor retained only 0.002% viability (Fig. S2). Overall, the above survey points to an industry consolidated around a small number of gram-positive microorganisms that are easy to stabilize and poor options for gram-negative organisms like *E. coli* Nissle 1917. In contrast, the synthetic biology and microbial therapeutics research fields in academia are consolidated around gram-negative bacteria with many of the most-forward looking examples (e.g. cancer treatment) using specifically *E. coli* Nissle 1917 ^2,9,10^.

### High throughput pipeline for generating synthetic extremophiles

To overcome the incongruity between the commercial and academic fields, we developed a high throughput pipeline to generate synthetic extremophiles (Fig. 2A, Methods). We defined a synthetic extremophile as a microbial formulation that has an enhanced capacity to survive insults such as drying, heating, and exposures to pressure, acid, organic solvents, and ionizing radiation. Much like natural extremophiles (e.g. spore-formers, tardigrades), such resilience would allow a synthetic extremophile to survive through the various manufacturing processes and long-term storage needs in the pharmaceutical, agricultural, and space health fields.

**Figure 2.**
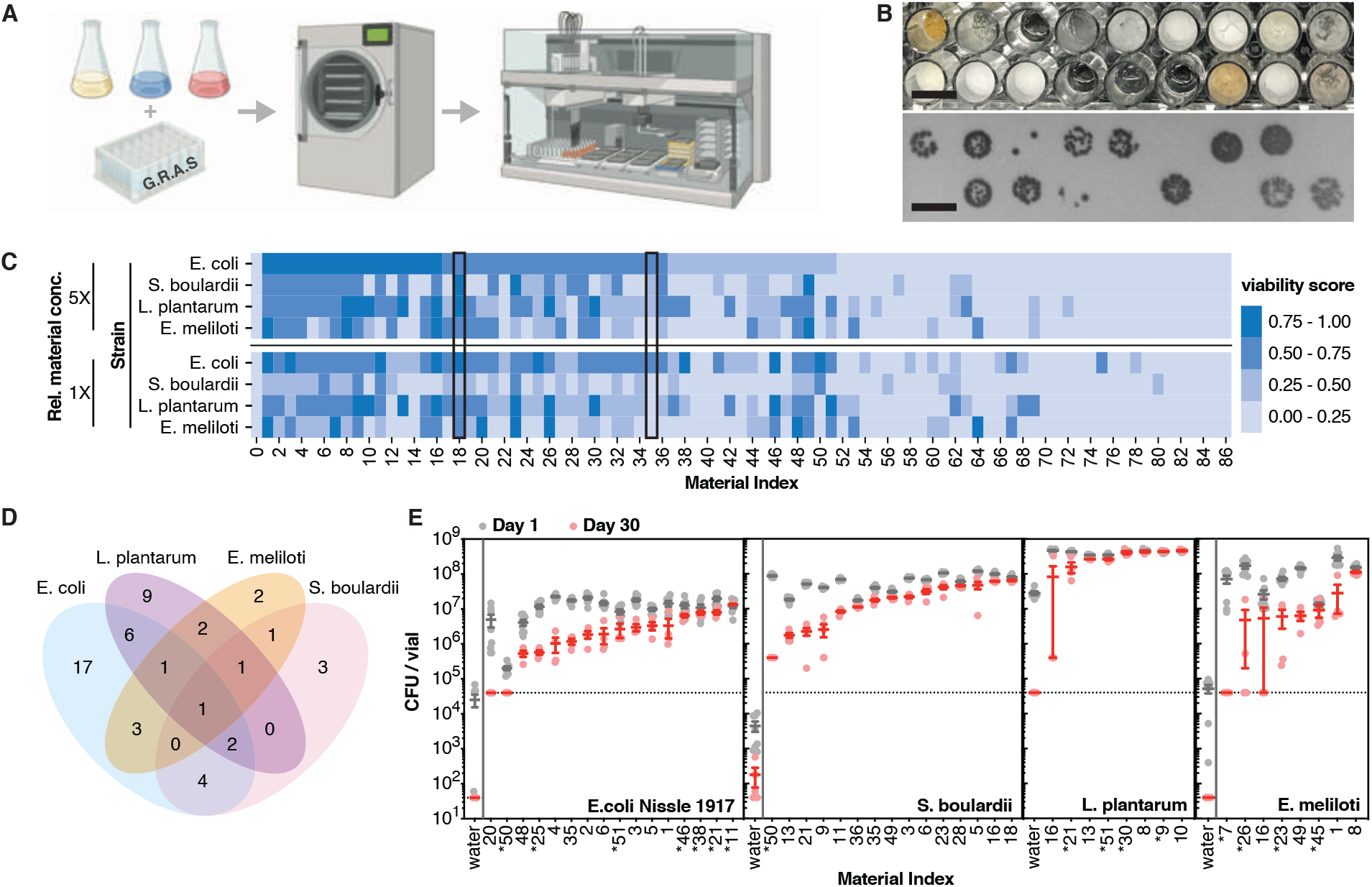
Development and validation of high throughput pipeline for dry stabilized microbial materials. **A**. We developed a high throughput pipeline (batch mixing, freeze-drying, plating and quantification) to assess the capacity of a library of materials generally recognized as safe (G.R.A.S) to stabilize microorganisms (see Methods). Illustration created with BioRender.com. **B**. Representative dried formulations made in high through put and corresponding colony counting images. Scale bars = 1 cm. **C**. Result of top 33% materials showing increased viability after drying across 4 organism and 2 concentrations and storage for 24 hours at room temperature, ordered by descending capacity to stabilize *E. coli* Nissle 1917 at the 5X concentration. The viability score is a composite colony count normalized to the maximum observable growth defined as 1 (see Methods). See Table S2 for material identities and concentrations corresponding to the noted material indices and Fig. S4 for full results. **D**. Correlation analysis of top performing materials for each organism. **E**. Verification of top materials defined in panel C via a more precise assay (freeze drying in vials and individual CFU counts on plates) and the ability to preserve viability for 1 and 30 days at room temperature. Stared material indices are at 1X concentration, otherwise at 5X. Horizontal dashed line represents the limit of detection. Where noted by a vertical bar the water sample was assayed at higher concentrations to achieve a lower limit of detection. N = 4 – 8 independently dried vials. Mean and SEM plotted over individual replicates.

We hypothesized that a synthetic extremophile that could survive in the dry state could then be enhanced to survive additional environmental insults using established material formulation techniques applied to the dry powder (e.g. coating)^11^. Therefore, we first screened a library composed primarily of materials generally recognized as safe (GRAS) by the Food and Drug Administration (FDA) for their ability to protect microbes through freeze-drying and survival at room temperature for 24 hours (Table S2). We applied this materials library to 4 microorganisms (*E. coli* Nissle 1917, *Ensifer meliloti, Saccharomyces boulardii, Lactobacillus plantarum*) that span a wide range of phylogenetic backgrounds, are technologically important and genetically tractable. Each microorganism was mixed with each of the 260 materials at two different concentrations to give 2,080 microbial-material formulations.

The residual viability of each formulation was measured via colony counts assessed in high throughput resulting in a normalized viability score (Fig. 2B, S3 and Methods). As expected, the included negative controls (i.e. antimicrobial compounds sodium metabisulfite and sodium hydroxide) consistently gave viability scores of 0 and the positive controls (i.e. known lyoprotectants trehalose and ATCC reagent 20) consistently resulted in measurable viability (Fig 2C and S4, black boxes). Additionally, as expected from the lyoprotectant literature^7^, sugars and peptones were overrepresented in top performing formulations (Fig. S5). Surprisingly, neither the positive controls nor any one single material was a top performer across the range of organisms. Instead, our results show that there are ideal species-specific formulations, with low overlap in best-performing materials even between the two gram-negative representatives (i.e. *E. coli* Nissle 1917 and *E. meliloti*) (Fig. 2D).

Next, we further stratified the top performing materials for each of the four microorganisms by evaluating colony counts after extended storage for one month at room temperature and relative humidity <5% (Fig. 2E). We chose these conditions to differentiate microbial-material combinations by their thermotolerance. This is in contrast to current microbial freeze-drying practice which still requires refrigeration to maintain acceptable viability in the dry state^7^. Stratifying in this way also provided a fingerprint for the intrinsic extremophilic qualities of the chosen organisms. For example, our results indicate that the lactic acid bacteria (*L. plantarum*) already had good thermotolerance as nearly all the top performing materials were indistinguishable after storage (Fig. 2E). Conversely, the two gram-negative bacteria showed a steeper drop-off with only a handful of materials maintaining the initial viability long term. These results further corroborate our probiotic viability survey suggesting that the lactic acid bacteria that dominate the market may be naturally more tolerant and easily stabilized by a wide range of materials.

### Optimization of the E. coli Nissle 1917 synthetic extremophile

Due to its importance in the microbial therapeutics and synthetic biology fields we next chose to further enhance the extremophilic qualities of *E. coli* Nissle 1917. Our screen showed that previously established mixtures, such as the positive control ATCC reagent 18, tended to outperform the individual constituent components also included in our screen (i.e. tryptic soy broth, sucrose, albumin). Therefore, we next compared two-material combinations of the library members with the best performing material (melibiose) against the single-material formulations and evaluated the residual viability after 24 hours both at room temperature as well as at 50 °C (Fig. 3A). We defined top hits as material combinations with viability scores above the control (melibiose + vehicle). From these we excluded library members that were themselves mixtures (e.g. LB broth) or of animal origin (e.g. mucin) to ease future manufacturing. The stability of the chosen two-material formulation (melibiose + fructooligosaccharides, caffeine or yeast extract) was verified in a larger format at 37 °C (Fig. S6). Of these, the mixtures with caffeine and yeast extract were robust and we next titrated their relative concentrations to find the optimal formulations which we termed formulation D and E (Fig. 2B).

**Figure 3.**
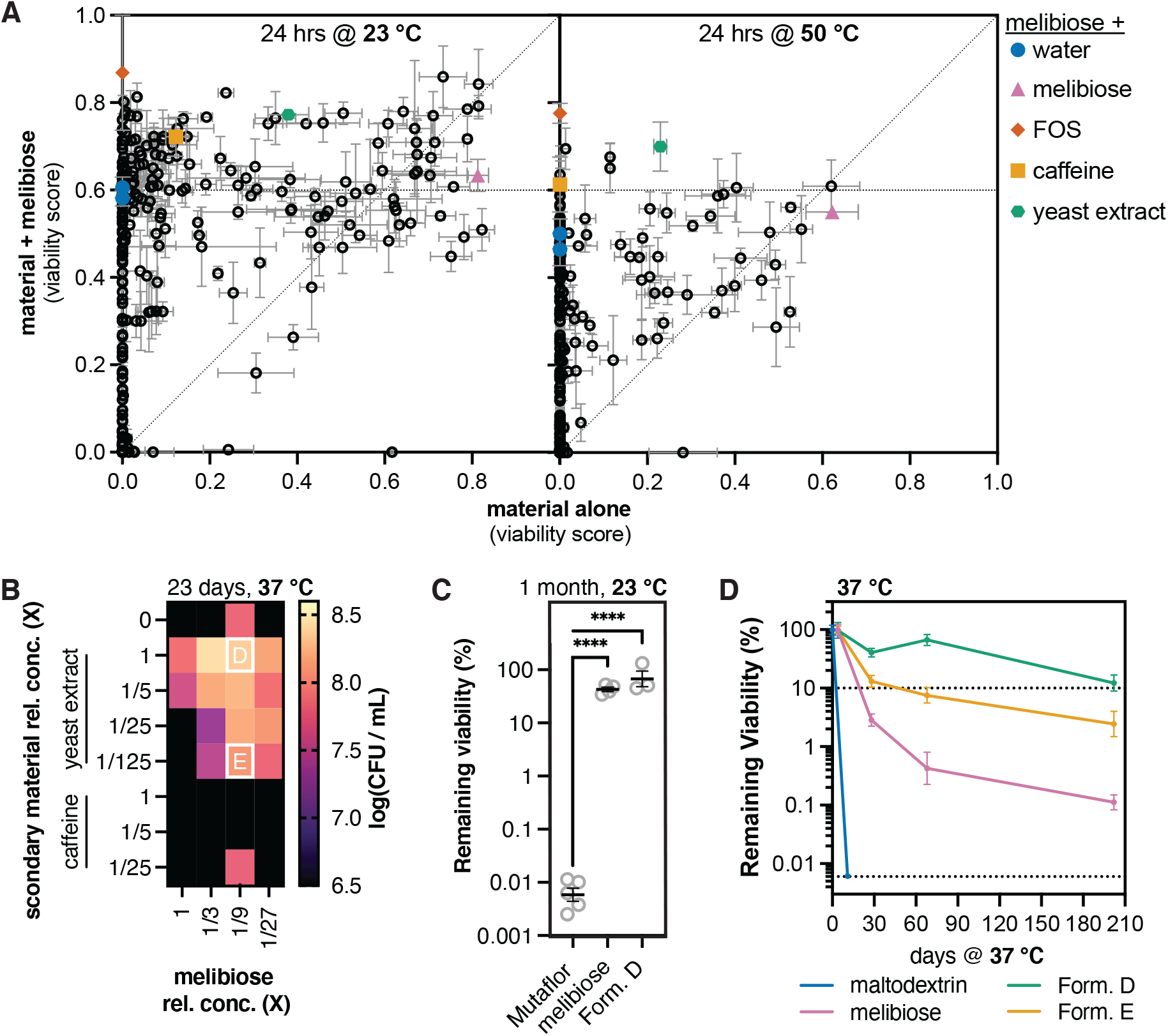
Application of pipeline to make *E. coli* Nissle 1917 into a synthetic extremophile. **A**. The library of materials was assayed individually (X-axis) and in combination with melibiose (Y-axis) for its capacity to stabilize *E. coli* Nissle 1917 at room temperature and 50 °C for 24 hours. Black circles mark the mean viability score (see Methods) along both dimensions and error bars represent the SEM. N = 3. Colored shapes mark material combinations with melibiose selected for further characterization. Horizontal dashed line is a reference that marks the viability score of melibiose combined with vehicle (water, blue circle). Diagonal dashed line is the identity. All concentrations were at 1X as defined in Table S2. **B**. The relative concentrations of each of the components in the selected two-material combinations (melibiose + yeast extract or caffeine) were varied, and the resulting effect on viability was measured after storage at 37 °C for 23 days. Concentration are all relative to 1X as defined in Table S2. White letters “D” and “E” mark the selected formulations for further characterization. **C**. Direct stability comparison of formulation “D” (as noted in panel B), the parent formulation (melibiose), and capsules of the commercial *E. coli* Nissle 1917 product Mutaflor after storage at room temperature for 1 month. Mean and SEM plotted over individual replicates. N = 3-5. Ordinary one-way ANOVA: ****, P < 0.0001. **D**. Characterized of ultra-long stability at 37 °C compared to maltodextrin (the stabilizer used in Mutaflor). Mean and SEM plotted for each timepoint. N = 3. Lower dashed line is the limit of detection.

Both melibiose and formulation D outperformed the commercial product Mutaflor (*E. coli* Nissle 1917) by over 3.5 orders of magnitude when stored at room temperature for 1 month (Fig. 2C). To confirm these results and exclude any manufacturing bias (i.e., freshly made vs store-bought product), we also formulated *E. coli* Nissle 1917 in maltodextrin (the stabilizer used in Mutaflor) using our specific freeze-drying processes. We then compared this “commercial-like” dry *E. coli* Nissle 1917 with our top performing formulations. We found that at 37 °C the maltodextrin stabilized material lost all measurable viability (>4 orders of magnitude) within 11 days (Fig. 2D). In contrast, formulation D retained over 10% viability even after 6.5 months at 37 °C (Fig. 2D and S7). This was >2 orders of magnitude better than the unoptimized melibiose formulation. In addition to maintenance of relative viability we also showed that the absolute CFU count could be optimized by altering the growth medium (Fig. S8), time of harvest (Fig. S9) and percent loading of bacteria (Fig. S10) to produce synthetic extremophiles with viabilities >10^10^ CFU per gram (Fig. S11).

### Compatibility of synthetic extremophiles with pharmaceutical manufacturing methods

These synthetic extremophiles open the field of potential use cases for microbial products to treat disease, enhance agricultural output and support exploration space travel (Fig. 4A). One immediate opportunity for human and animal health would be the ability make dosage forms with tuned release parameters (e.g. delayed release, enteric coatings, etc). However, this requires applying pharmaceutical manufacturing techniques normally too harsh for living cells such as milling (exposure to shear forces), wet granulation (exposure to isopropanol and baking), tableting (exposure to high pressure) and spray coating (exposure to acetone and baking). Therefore, we submitted the *E. coli* Nissle 1917 synthetic extremophile to this battery of processes (Fig. 4B) and characterized its robustness compared to the current commercial alternative (maltodextrin formulation) (Fig. 4C). We found that the synthetic extremophile could be milled, mixed with excipients (i.e. binder, glidant, filler) and tableted in open air at room temperature and still retain > 15% viability which was 2.4 orders of magnitude higher than the commercial control (0.06% viability) (Fig. 4C). Wet granulation was similarly well tolerated (>28% viability) and we found that tableting pressure modulates the ultimate shelf stability (Fig. S12).

**Figure 4.**
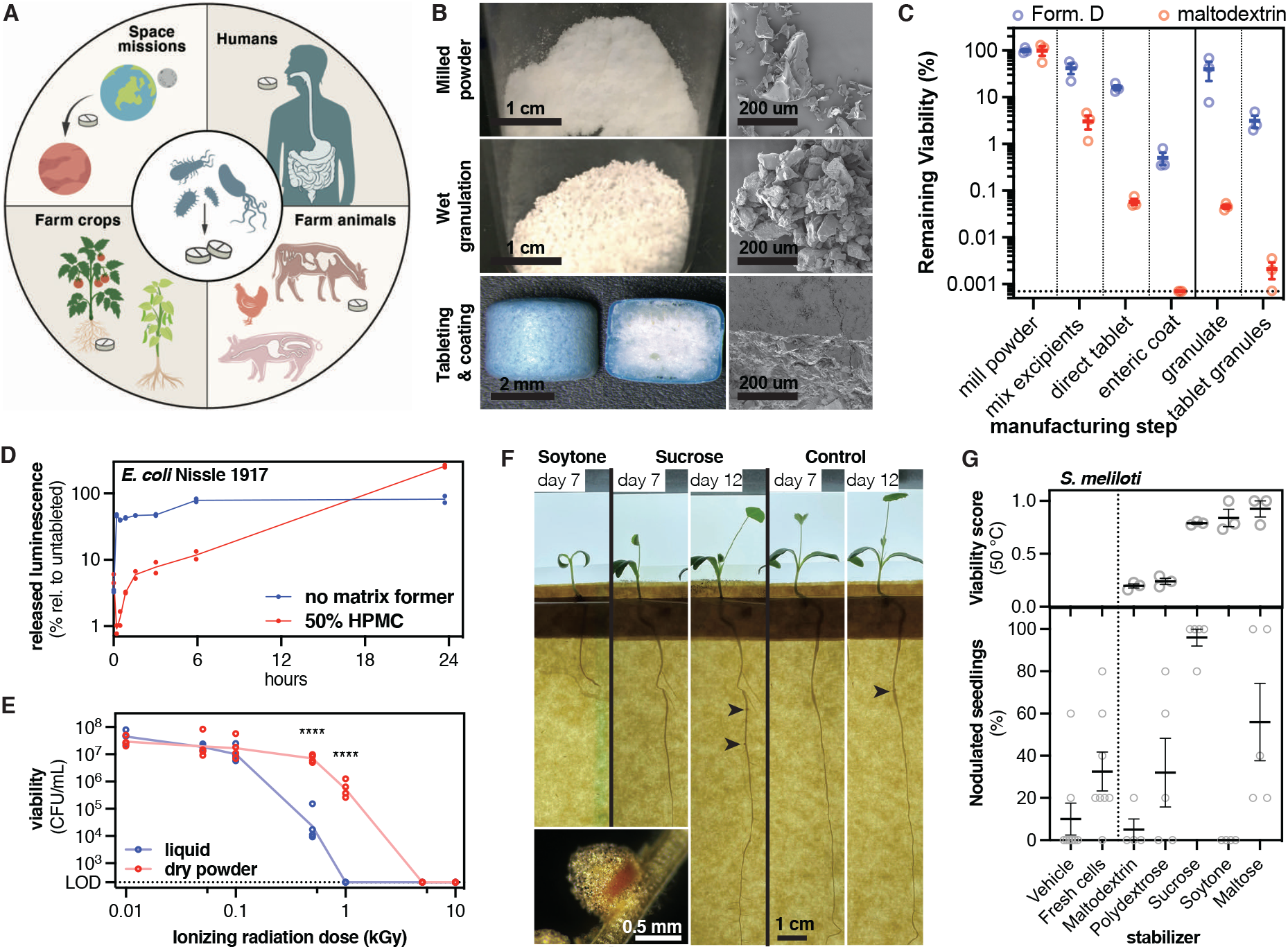
Synthetic extremophiles withstand harsh processing techniques to make microbial therapeutics for humans, agriculture, and space. **A**. Microbial tableting and coating allows simple dosing, transport and tuning of release parameters to access a wide range of applications. **B**. Representative photos and scanning electron micrographs of synthetic extremophiles process through the industrially relevant processes of milling, wet granulation, tableting and spray coating. **C**. Stability of the *E. coli* Nissle 1917 synthetic extremophile (melibiose) through each sequential manufacturing process relative to the parent powder. Maltodextrin is the stabilizer used in the commercial product Mutaflor. Horizontal dotted line marks the limit of detection. Mean and SEM plotted over individual replicates. N = 3. **D**. Tableting allows tunning of release kinetics through the inclusion of a matrix former (hydroxypropyl methylcellulose, HPMC). Synthetic extremophile *E. coli* Nissle 1917 cells carry a biosynthetic pathway (luxCDABE plasmid) and the release luminescence was tracked over time. Lines connect the means. Individual replicates plotted. N = 2. **E**. High viability in the dry state allows higher resistance to ionizing radiation. Synthetic extremophile *E. coli* Nissle 1917 cells or a liquid suspension of the cells in PBS were exposed to the noted ionizing radiation dose. Lines connect the means. Individual replicates plotted. N = 4. Two-way ANOVA: ****, P < 0.0001. **F**. *E. meliloti* is a nitrogen-fixing bacteria that supplies plants with nitrogen at symbiotic root nodules (inset). Synthetic extremophile *E. meliloti* where exposed to 50 °C, hydrated and tested for functionality in a plant root nodulation assay with *M. truncatula*. Nodulated seedlings (black arrows) can be quantified at day 12 post root inoculation. **G**. Quantification of nodulation assay (panel F) compared to the material viability score. Mean and SEM plotted over individual replicates. N = 3 - 5 and 8 for controls (left of dashed line).

Once tableted, the synthetic extremophile also survived spray coating with an acetone solution of an enteric polymer (Eudragit L100-55) whereas the commercial alternative lost all viability (Fig. 2C). In this state not previously accessible by *E. coli* Nissel 1917, we found that the synthetic extremophile could retain 100% viability through an exposure to simulated gastric fluid at pH 1.2 for 1 hour at 37 °C (Fig. S13).

### Compatibility of synthetic extremophiles with ionizing radiation

Next, to validate the compatibility of the synthetic extremophile with use in exploration space flight, we characterized its hardiness to a range of ionizing radiation doses from 100 to 10,000 Gy (Typical radiation dose values: ∼15 uGy/day on Earth^12^, ∼200uGy/day on the Moon^13^, ∼250 uGy/day on Mars^14^, total accumulated dose 500 day trip to Mars is 0.32 Gy^15^, 444 days outside the ISS is 0.41 Gy^16^). The maximum value of 10 kGy represents the dose generally accepted for radiation-based sterilization of food.^17^ We found that the synthetic extremophile could sustain exposures up to 1000 Gy (Fig. 4E). At this radiation level a liquid suspension of the same bacteria lost all measurable viability.

### Functional characterization of synthetic extremophiles

We next asked whether the synthetic extremophile could also remain functional in the tableted state and whether that function could be tuned through dosage form design. To test this, we incorporated a bioluminescent biosynthetic pathway (luxCDABE) measured luminescence upon rehydration. While the dry tablet showed no measurable luminescence, 48% of the ultimate luminescent level was reached within 15 minutes of rehydration (Fig. 4D). Substituting the tablet filler by the matrix forming material hydroxypropyl methylcellulose (HPMC) lead to a slow release of the bacteria and tuning of the kinetics of the luminescent function over a period of 24 hours (Fig. 4D). Combinations of immediate and slow-release tablet layers can be used to make a mixed microbial pill that seed different bacteria in the host microbiome at different rates.

Finally, due to its potential for increasing the sustainability of agricultural systems with a lower need for chemical fertilizers, we asked whether our approach to making the synthetic extremophiles would interfere with the complex biological process of symbiotic nitrogen fixation in plant roots^3,18^. This symbiotic process occurs between legumes and bacteria (rhizobia) in the soil, via the formation of root nodules which house the bacteria that can then fix atmospheric nitrogen (N2) into a usable form for the plant^18^. Specifically, we chose to evaluate nodule formation by the nitrogen-fixing bacteria *E. meliloti* in *Medicago truncatula*, a model plant used for studying this process^18,19^.

*E. meliloti* is currently used as a seed inoculant prior to planting, however, there remains a need for improved stability of the microbial product during the adverse conditions of seed inoculation and storage, and in particular tolerance to elevated temperatures above 40 °C^20,21^. Therefore, to generate a synthetic extremophile version of *E. meliloti*, we repeated the heat shock screen at 50°C to find materials that would give *E. meliloti* synthetic thermotolerance (Fig. S14). From these we chose three optimal materials (sucrose, soytone, maltose) and two suboptimal materials (maltodextrin, polydextrose). *E. meliloti* was formulated in each, submitted to a 50 °C shock for 24 hours, rehydrated and applied to seedlings of *M. truncatula* A17 (Fig. 4F). Functional symbiosis was measured as percent of seedlings that successfully formed nodules. The synthetic *E. meliloti* extremophile made with sucrose and maltose were able to successfully nodulate while the one made from soytone lead to plant stunting (Fig. 4F and G). These results show that our high throughput pipeline can define a set of materials that are not just ideal for the target microbial species, but which can be further selected for those that are compatible with the specific target application that may include complex biological processes such as nodulation.

In sum, we present an approach for the identification of materials capable of imparting microorganisms with extreme environment tolerance. These synthetic extremophiles stand to transform our capacity to disseminate bioactive organisms across human applications, from shelves across the globe, to fields for agricultural practices to space exploration.

## Supplementary Materials

The methods, supplementary figures and supplementary tables are available as a separate file.

## Acknowledgments

We thank R. Langer for providing guidance, feedback and experimental facilities. We thank Anna Hupalowska for creating the illustration in Figure 4A.

## Author contributions

MJ, GT conceived and designed the research; MJ designed the high throughput pipelines and wrote scripts for liquid handling and image analysis; JL, ZV, AKS, QC established the calibration curves for microbial quantification; MJ, JL designed and performed the commercial probiotics survey; MJ, JL performed the initial material stabilizer evaluations and validations; MJ, EK performed the material stabilizer optimization and validations; MJ, EK designed and performed the synthetic extremophile pharmaceutical manufacturing and viability evaluation. MJ, JB designed and performed the ionizing radiation experiment; MJ, JL, KM designed, performed and optimized the plant nodulation assay; BM performed scanning electron microscopy of the microbial materials; MJ, JL, EK performed formal analysis of the data; MJ, GT wrote the manuscript; MJ, JL, EK, GT edited the manuscript; MJ, GT supervised and managed project progress and personnel; MJ, GT supervised and managed funding of the project.

## Data and materials availability

All data needed to evaluate the conclusions in the paper are present in the paper and/or the Supplementary Materials. Additional data related to this paper may be requested from the authors.

