## Supplementary material for "Synthetic extremophiles: Species-specific formulations for microbial therapeutics and beyond": Methods, and Supplementary Figures and Tables

**This PDF file includes:**

Materials and Methods
Supplementary references
Figs. S1 to S14
Tables S1 to S4

### Materials and Methods

Viability survey of commercial probiotic and microbial products (Figures 1, S1)

All products were purchased and analyzed in 2019 well before their expiration dates. The specific product names, lot numbers, expiration dates, date of analysis and dosage forms are summarized in Table S1. All procedures were analyzed under sterile conditions. The dosage forms encountered where two-piece capsules filled with loose powder, sachets (packets) filled with loose powder or tablets. In each case only the microbial fraction was analyzed. Specifically, for capsules, the two halves were carefully separated, and the internal contents poured out onto weighing paper for further analysis. For sachets, the sachets were carefully torn open, and the internal contents poured out onto weighing paper for further analysis. For tablets, the tablet was placed in a sterile zip-loc bag, crushed by manually rolling a plastic tube over the tablet, and the resulting powder was poured out onto weighing paper for further analysis.

*Promised colony forming unit (CFU) counts per dose* were recorded from the product packaging and converted to CFU per gram by dividing by the mean mass of 4 doses. For Florastor, the amount of yeast cells was reported in milligrams per dose, and this number was converted to CFU per dose by dividing by the approximate cell dry weight of yeast (~20 pg, 32% of 60 pg)[BNID 101795, BNID 105094][1]. The dose mass only includes the recovered microbial fraction as described above (i.e. internal contents of the dosage form). In cases that products report a CFU count both "at manufacture" and "by expiration date" only the CFU count "by expiration date" was used. This data is summarized in Table S1.

*Viable CFU counts per dose* were determined by rehydrating one dose in phosphate buffered saline (PBS, ThermoFisher 10010049) at the specified rehydration ratio on ice, making 10-fold serial dilutions in PBS on ice, plating 100uL of each dilution onto the appropriate solid medium in 100 mm circle petri dishes with 4.5mm plating beads (Zymo), incubating at the appropriate conditions, and counting colonies on the dilution that gave ~200-1000 distinct colonies. Four or five separate doses of each product were prepared as described. As noted in Table S1, the rehydration ratio was typically one dose in 50 mL, but in a few instances the rehydration volume was lowered when the cell counts were found to be low during an initial analysis. The medium and culture conditions used for each product were chosen based on the microorganisms listed on the package and are listed in Table S1. Medium components were all purchased from BD Life Sciences as specified in Table S3, solidified with 1.5% w/v Bacto Agar (BD, 214010) and prepared according to the label instructions. For aerobic conditions plates were incubated in a static incubator and for anaerobic conditions plates were first placed in a BD GasPak EZ Gas Generating Systems container (BD, 260002) with an anaerobe sachet (BD, 260001) and incubated in a static incubator. Colony counts were determined systematically by first imaging the plates in a gel imager (ChemiDoc XRS+, BioRad) using UV transillumination and the standard emission filter (580/120), and then programmatically counting the colonies using FIJI (imageJ)[2]. Total viable CFU counts per dose were calculated by multiplying the colony counts by the plated dilution factor (to get a CFU / mL value) and then the total rehydration volume. Viable CFU counts per dose where then converted to CFU counts per gram by dividing by the mass of the specific dose analyzed. A dose only includes the recovered microbial fraction as described above (i.e. internal contents of the dosage form).

*Total cell counts per dose* were determined using an automated microscopic cytometer (QUANTOM Tx Microbial Cell Counter, Logos Biosystems) according to the manufacturer protocols. Specifically, a 10 uL sample of the rehydrated dose used for the viable CFU counts (above) was mixed with 2 uL of a 1:1 mix of total cell staining dye and enhancer (Logos

Biosystems, Q13501) on ice, then 8 uL of cell loading buffer (Logos Biosystems, Q13501) and 6 uL of the mixture were loaded onto hemocytometer slides (Logos Biosystems, Q12001). Slides were centrifuged for 10 minutes at 300 rcf in a slide microcentrifuge (Logos Biosystems, Q10002) and then loaded into the cell counter. Four or five separate doses of each product were prepared as described, and each prepared sample was loaded and quantified in duplicate. For products that gave counts above 1e9 cells per mL during an initial analysis, the rehydrated dose was first diluted 10-fold in PBS before sample preparation. Total cells per dose were calculated by multiplying the cells per mL as reported by the instrument by the dilution factor (if any) and the total rehydration volume as specified in Table S1. Total cell counts per dose where converted to cell counts per gram by dividing by the mass of the dose analyzed. A dose only includes the recovered microbial fraction as described above (i.e. internal contents of the dosage form).

Percent viable cells relative to promised cells were calculated by dividing the viable CFU counts per gram by the promised cell counts per gram. Percent viable cells relative to total cells were calculated by dividing the total CFU counts per gram by the total cell counts per gram.

Heat stress test of commercial probiotic and microbial products (Figure S2)

Three or four doses of each product as listed in Table S1 were incubated at 50 °C for 24 hours in their original dosage form in a static incubator. Three or four control doses were kept at the manufacturer recommended storage temperature (4 °C or 23 °C as noted in Fig. S2) for the same period. The viable CFU counts per gram for heat stressed and control doses were determined as described above ("Viability survey of commercial probiotic and microbial products"). Percent retention of viability was calculated by dividing each heat stressed CFU count per gram by the mean of the control CFU counts per gram.

Microbial strains used for stabilization and general culture conditions

Details of microbial strains used are listed in Table S4 and medium components are listed in Table S3.

*E. coli* Nissle 1917 was isolated from the commercial product Mutaflor and routinely cultured on LB agar at 37 °C in a static incubator or LB broth in 14 mL culture tubes or vented, baffled culture flasks shaken at 250 rpm at 37 °C in a shaker incubator.

*S. boulardii* was isolated from the commercial product Florastor and routinely cultured on YPD agar at 30 °C in a static incubator or YPD broth in 14 mL culture tubes or vented, baffled culture flasks shaken at 250 rpm at 30 °C in a shaker incubator.

*E. meliloti* Rm1021 was purchased from the American Type Culture Collection (ATCC) and routinely cultured on TY agar at 30 °C in a static incubator or TY broth in 14 mL culture tubes or vented, baffled culture flasks shaken at 250 rpm at 30 °C in a shaker incubator.

*L. plantarum* NC8 was purchased from the Culture Collection University of Gothenburg (CCUG) and routinely cultured on MRS agar or MRS broth in sealed 14 mL culture tubes or sealed culture flasks at 37 °C in a static incubator.

High throughput pipeline for microbial material stabilizers (Figures 2ABC, 3A, S3, S4, 4G, S14)

*Microbial strains* were precultured overnight in 5 mL liquid medium at the appropriate conditions (see above). The overnight culture was diluted 1:1000 fold into three replicates of 125 mL of fresh liquid medium in a flask and cultured for 24 hours. The optical density ($OD_{600}$) of each replicate culture was recorded (typically 2 – 2.5 for *E. coli* Nissle 1917, *E. meliloti, L. plantarum* and 5 – 6 for *S. boulardii*) and the cells were pelleted at 3220 rcf for 15 minutes. The cell pellets were then resuspended in PBS (*E. meliloti, L. plantarum*, *S. boulardii*) or spent medium (*E. coli* Nissle 1917) to a final $OD_{600}$ of 2.25 and used immediately.

*The material library* was prepared by mixing each material with ultrapure water to the specified concentration (1X, 5X) in Table S2. The vendors and product numbers for all materials are listed in Table S2. The material solutions were arrayed in sealed deep well plates and kept at -20 °C until needed. On the day of use, the material plates were thawed at room temperature, vortexed well and centrifuged briefly.

*For the two-material combinations with melibiose*, the arrayed material library was first mixed 1:1 with melibiose or water as a control at the 1X concentration of all components (Table S2). This gives a 0.5X final concentration of each component in Figure 3A relative to the one-material library results reported in Figures 2C and S4.

*To freeze-dry*, each replicate microbial cell suspension (25 uL) was mixed with the arrayed materials (75 uL) in batch into flat-bottom 96-well plates using a liquid handling robot (Tecan, EVO 150). The arrayed microbial-material plates were immediately placed into a tray freeze dryer (Labconco, FreeZone Stoppering Tray Dryer) with shelfs pre-cooled to -40 °C and on custom 0.25-inch aluminum pedestals to ensure heat transfer to the plates. The samples were annealed at -20 °C and then dried at -20 °C and 0.1 mBar for 12 hours (nominal) and then at 37 °C for 3 hours (nominal). The chamber was backfilled with nitrogen, opened to atmosphere and then the plates were covered loosely with parafilm, capped, and placed in nitrogen-flushed zip-loc bags with desiccant (Drierite, granular -8 mesh). The bagged, freeze-dried plates were stored in the dark at the specified temperature (23 °C or 50 °C) for 24 hours.

*The viability* of the stored microbial-material combinations was determined by rehydrating in water, making 10-fold serial dilutions in PBS, plating onto the appropriate solid medium and quantifying the resulting colonies. Specifically, using a liquid handling robot (Tecan, EVO 150), each replicate plate was rehydrated with ultrapure water (200 uL per well), diluted in PBS to make $1{:}10^2$, $1{:}10^3$, $1{:}10^4$ dilutions (relative to the initial volume before drying) and 4 uL of each of the dilutions were spotted onto solid medium in 1-well rectangular plates. After incubation at the appropriate culture conditions (see above) the plates were imaged in a gel imager (ChemiDoc XRS+, BioRad) using UV transillumination and the standard emission filter (580/120).

*To assign a Viability Score*, using FIJI (imageJ), the 16-bit tiff plate images were programmatically separated into individual spot images, each spot was segmented, and several image attributes were measured (raw particle count, mean background value, mean segmented region value, segmented region area). These were used to classify spot images into empty, countable and lawns using the mean segmented region value to background value ratio and segmented region area attributes (Figure S3). Raw particle counts for the empty class were set to 0. Raw particle counts for the countable class was interpreted as a true colony count. Raw particle counts for the lawn class were set to a microbe-specific colony count value determined by extrapolating the data observed on a raw count vs segmented region area plot (Figure S3). For *E. coli* Nissle 1917 and *E. meliloti* lawns were set to 60 counts. For *S. boulardii* lawns were set to 50 counts. For *L. plantarum* lawns were set to 120 counts. These values are sensitive to the colony size at the time of analysis. Finally for each microbial-material combination we calculate a Viability Score which is the cumulative sum of colony counts at the three dilutions divided by the maximum total possible count (i.e. all lawns). The Viability score thus ranges from 0 to 1. This method allows us to use the data across all three analyzed dilutions in order to average out the sampling noise caused from plating and counting low numbers of colonies (i.e. the three dilutions act as technical replicates). It also allows us to be agnostic to the actual number of countable colonies on any dilution and to assign a robust value to microbial-material combinations that lead to lawns in some dilutions and biological replicates (i.e. can assign a value to a wider range of viabilities that might be possible by using only the countable spots). This analysis was applied to three independent biological replicates for each material-microbial combination.

*Correlation analysis* of top performing materials for each organism was carried out by first normalizing the viability scores of each specific material and concentration to the maximum viability score observed for each organism. This gives a list of materials (at specific concentrations) for each organism with normalized scores that span the full range 0 to 1. Then any material with a normalized score above 0.75 was defined as a top material for that organism. Based on which organism sets each of these top performing materials (at specific concentrations) belonged to, they were binned into the 15 possible sets of 1-, 2-, 3- and 4-organism overlaps.

Vial-based viability validation of microbial material stabilizers (Figure 2E)

Top performing microbial-material combinations in the high throughput pipeline were assayed at a larger scale to determine a more accurate relative viability value (CFU / vial) and assess their short term (1 day) and long term (30 day) stabilization potential at room temperature. The analysis described below was repeated for 7 – 8 replicates for the 1 day time point and for 4 – 5 replicates for the 30 day time point.

*Microbial strains* were precultured overnight in 5 mL liquid medium at the appropriate conditions (see above). The overnight culture was diluted to an $OD_{600}$ of 0.00225 in 25 mL of liquid medium in a flask and cultured for 24 hours at the appropriate conditions (see above). The cells were harvested by centrifugation and resuspended in cold PBS at a final $OD_{600}$ of 2.25 and used immediately.

*To freeze-dry*, each microbial cell suspension (100 uL) was mixed with the target material (300 uL) with a micropipette in autoclaved threaded tube vials (Electron Microscopy Sciences, 60304-04) capped with 2-prong lyophilization stoppers (Electron Microscopy Sciences, 60304-41) to the first stop. Within 10 minutes of mixing, the vials were placed in a tray freeze dryer with shelfs pre-cooled to -40 °C and in aluminum StableTemp vial blocks (Cole-Parmer, EW-36600-44). Samples were dried as described above (see "High throughput pipeline"). After drying, the chamber was backfilled with nitrogen, opened to atmosphere, vials fully stoppered, sealed with screw caps, and stored in the dark at 23 °C for the specified time.

*The viability* of the stored samples was assessed by hydrating the samples with 4 mL of cold PBS (to give a 1:10 dilution relative to the original volume), making 10-fold serial dilutions, plating 100uL of each dilution onto the appropriate solid medium in 100 mm circle petri dishes with 4.5mm plating beads (Zymo), incubating at the appropriate conditions, and counting colonies on the dilution that gave ~200-1000 distinct colonies. CFU counts per vial were determined systematically as described above using a gel imager and FIJI (see "Viability survey of commercial probiotic").

Validation of two-material combinations (Figure S6)

*E. coli* Nissle 1917 was prepared as described above (see "Vial-based viability validation") with a few modifications: the flask culture was 2 L and incubated for 17 – 18 hours and the final cell suspension in PBS was set to an $OD_{600}$ of 10 – 12.

*To freeze-dry,* the cell suspension (12.5 mL) was mixed with the target material combination (37.5 mL), poured into rectangular 1-well plates, and placed into a tray freeze dryer with shelfs pre-cooled to -40 °C and on custom 0.25-inch aluminum pedestals to ensure heat transfer to the plates. Samples were dried as described above (see “High throughput pipeline”). After drying, the chamber was backfilled with nitrogen, opened to atmosphere, plates were transferred to zip-loc bags, the dry material was scrapped into the bag, milled by rolling a plastic tube on the outside of the bag and the resulting powder was kept in the bag with a desiccant pack and stored in the dark at 4 °C until needed.

*To prepare samples “with excipients”*, the dried bacterial powders were mixed with 1 % w/w magnesium stearate, 5% w/w polyvinylpyrrolidone and 64% w/w lactose to give a 30% w/w loading of the bacterial material.

*Long-term storage of samples* was done by placing ~20 mg (for bacterial powders alone) or ~60 mg (for bacterial powders with excipients) aliquots in 12-well plates, capping the plates and placing them in nitrogen-flushed zip-loc bags with desiccant (Drierite, granular -8 mesh). Bagged plates were stored at 37 °C in the dark for the indicated time.

*The viability* of the stored samples was assessed by hydrating the samples with 1 mL PBS, making 10-fold serial dilutions, spotting ~1.5 uL of each dilution in triplicate onto LB agar in 1-well rectangular plates using a pin replicator (V&P Scientific, VP 407A), incubating at 37 °C, and counting colonies on the strongest dilution that gives > 1 colony per spot. Total CFU counts per sample were calculated by first dividing spot counts by the spotted volume and multiplying by the dilution factor and total rehydration volume, and then averaging across the three spotting replicates. Total CFU counts per mass were calculated by dividing the CFU counts per sample by the mass of the stored sample. Three independently stored replicate samples were analyzed as described. The percent viability at each time point was calculated by dividing the total CFU counts per mass by the mean total CFU counts per mass of three replicate samples at day 0.

Note: this method tends to systematically underestimate the true viable CFU counts but is a robust measure of relative viability.

<u>Optimization of two-material formulations (Figure 2B)</u>

To find optimal concentrations for each of the components in the selected two-material formulations for *E. coli* Nissle 1917, we screened a combinatorial library of component concentrations using the high throughput pipeline described above (see “High throughput pipeline for microbial material stabilizers”) with a few modifications: the flask culture was 2 L and incubated for 17 – 18 hours, the final cell suspension in PBS was set to an $OD_{600}$ of 10 – 12, dried plates were stored at 37 °C for 23 days, viability was assessed at the $1:10^5$ dilution where only empty or countable spots were present and the raw counts were converted directly to CFU / mL by dividing the spot counts by the spotted volume (4 uL) and multiplying by the dilution factor.

The top performing combinations were named Formulation D (1/9X melibiose + 1X yeast extract) and Formulation E (1/9X melibiose + 1/125X yeast extract).

<u>Direct comparison of synthetic extremophile *E. coli* Nissle 1917 with Mutaflor (Figure 2C, S11)</u>

Stabilized *E. coli* Nissle 1917 was prepared as described above (see “Validation of two-material combinations) with either melibiose (1X) or Formulation D (see “Optimization of two-material formulations”). For 100X cultures the 2 L culture was concentrated 100X in PBS before mixing with the stabilizers. Mutaflor capsules were kept intact as manufactured during storage.

*Samples were stored* in nitrogen-flushed zip-loc bags, with desiccant (Drierite, granular -8 mesh), kept at 23 °C in the dark for the indicated time.

*The viability* of the stored samples was determined as described above (see “Viability survey of commercial probiotic and commercial products”).

Ultra-long-term assessment of viability (Figure 2D, S7)

Stabilized *E. coli* Nissle 1917 was prepared as described above (see “Validation of two-material combinations) with either melibiose, Formulation D, Formulation E or 5 % w/w maltodextrin (16.5-19.5 DE) as the commercial control. It was determined from the list of inactive ingredients that maltodextrin was the only stabilizer used in the *E. coli* Nissle 1917 product Mutaflor.

Multi-gram samples of the dry powders were stored in 20 mL glass with desiccant and kept in the dark at the specified temperature. Viability was assessed by retrieving a small quantity (~60 mg) of the stored powders and following the procedure described above (see “Validation of two-material combinations”) but the samples were rehydrated in ultrapure water instead of PBS. In addition to the pin replicator spotting, one sample was also traditionally plated onto 100 mm circle petri dishes as described above (see “Validation of two-material combinations”) to determine the scaling factor between the spotting method and the petri dish plating. When CFU / mg is reported we use the scaled values. Percent viability is not impacted by this scaling factor as it is a relative value.

Viability after exposure to wet granulation, tableting and enteric coating (Figure 4BC)

Stabilized *E. coli* Nissle 1917 was prepared as described above (see “Validation of two-material combinations) with either Formulation D or 5 % w/w maltodextrin (16.5-19.5 DE) as the commercial control. Dry powders were mixed with the following excipients: 1 % w/w magnesium stearate, 5% w/w polyvinylpyrrolidone and 64% w/w lactose (this gives a 30% w/w loading of the bacterial material).

*Wet granulation* was carried out by adding 600 uL isopropanol to 2 g of the bacterial-excipient mixture while it was being continually stirred with a stand mixer (Sunbeam, B000COC69C) room temperature. Stirring was allowed to continue for an additional 5 minutes, and the resulting paste was passed through a bench top oscillating granulator (ERWEKA, FGS II) with a 1 mm mesh screen. The collected granules were dried at 45 °C for 20 min in a food dehydrator (Magic Mill, MFD-9100) retrofit with a HEPA filter (Vornado, MD1-0022) to maintain sterility. Granules were stored in tubes with desiccant at 4 °C until used.

*Tableting* was carried out in the open air on a tablet press (Natoli, NP-RD10A) with a 5 x 5 mm circular punch and die, set to 8 mm depth, and pressed with 17 kN of force (3.4 kN per tablet equal to a pressure of 173 uPa). Either the bacterial-excipient mixture was tableted directly (“direct tablet”) or the granulated mixture was tableted (“tableted granules”). Tablets were stored in tubes with desiccant at 4 °C until used.

*Enteric coating* of the tablets was done by spray coating with a solution of 3.6 % w/w of Eudragit S100 (Evonik) in a 1:1 co-solvent of acetone and isopropanol with 0.36 % w/w triethyl citrate as a plasticizer. Spray coating was carried at 23 °C via a spray nozzle with a 0.5 mm opening with the tablets in a rotating pan coater (ERWEKA, DKE) over the course of 1 hour. Coated tablets were dried at 40 °C for 2 hours in a food dehydrator as described above. Coated tablets were stored in tubes with desiccant at 4 °C until used.

*The viability* of each of the processed samples was assessed as described above (see “Validation of two-material combinations”) but samples were rehydrated in water instead of PBS, and the CFU per mass was calculated by dividing my the mass of only the bacterial fraction (i.e. 30% of total mass for samples that include excipients). Coated tablets were first cut in half to expose the center before rehydration. Three independent samples of each type were assessed as described.

Viability after exposure to simulated gastric fluid (Figure S13)

Enteric coated tablets of *E. coli* Nissle 1917 were prepared as described above (see "Enteric coating") were placed in 40 mL of simulated gastric fluid (USP, pH 1.2, no enzymes) in a 50 mL conical bottom tubes and incubated for 1 hour at 37 °C on an oscillating tube revolver (Thermo Scientific, 88881001) inside a static incubator. After exposure to simulated gastric fluid, tablets were blotted dry, cut in half and the viability was assessed as described above (see "Validation of two-material combinations").

Viability after exposure to ionizing radiation (Figure 4E).

Stabilized *E. coli* Nissle 1917 was prepared as described above (see "High throughput pipeline for microbial material stabilizers") with a few modifications: only melibiose (1x) was used as the stabilizer, 32 replicate samples were arrayed on eight 96-well plates (4 replicates per plate) and bagged plates were stored at 4 °C until used. On the day of irradiation, a fresh overnight 5 mL culture of *E. coli* Nissle 1917 was arrayed onto the same plates (4 replicates per plate) as a control.

The plates were sequentially placed into a Gammacell irradiator (Best Theratronics, 40 Exactor) and exposed to a Cobalt-60 source for the time required to achieve the specified ionizing radiation dose (dose rate = 37.12 Gy/min). One plate was maintained with the other plates but not exposed to any radiation.

The viability of each of the samples was assessed as described above (see "Validation of two-material combinations") except that samples were rehydrated with 200 uL of PBS directly in the treatment plates to give an initial 1:2 dilution for subsequent serial dilution.

Tuning of release kinetics via matrix formers (Figure 4D)

*E. coli* Nissle 1917 was transformed with a plasmid encoding a constitutively expressed pathway for bioluminescence (luxCDABE) via electroporation. Plasmid and strain details are listed in Table S4.

Luminescent *E. coli* Nissle 1917 stabilized with Formulation D was prepared as described above (see "Validation of two-material combinations) with cultures supplemented with 100 ug/mL of ampicillin. The resulting bacterial powder was mixed either with the standard excipients (1 % w/w magnesium stearate, 5% w/w polyvinylpyrrolidone and 64% w/w lactose) or with the inclusion of a matrix former (50 % w/w hydroxypropyl methylcellulose, 1 % w/w magnesium stearate, 5% w/w polyvinylpyrrolidone and 14% w/w lactose). In both cases the bacterial material maintains a 30% w/w loading.

A fraction of each of these two powder mixtures was made into tablets as described above for the "direct tablet" method. Then either one tablet or the equivalent mass of loose powder (with excipients) was placed into 30 mL of ultrapure water in a conical bottom tube and incubated for 24 hours at 37 °C on an oscillating tube revolver (Thermo Scientific, 88881001) inside a static incubator. At the specified timepoints 200 uL samples were taken, arrayed in white-walled, clear-bottom 96-well plates and the luminescence quantified on a luminometer (Tecan, Infinite 200). Each of the four sample types (standard or matrix forming excipients in loose or tableted form) were analyzed as described in duplicate.

To calculate the percent released luminescence, at each time point the luminescence value of the tableted samples was divided by the mean luminescence value of the corresponding loose powder samples.

Plant nodulation assay with stabilized *E. meliloti* (Figure 4FG)

*E. meliloti* cultures were prepared as described above (see "High throughput pipeline for microbial material stabilizers") and freeze-dried with the specified stabilizers following the 1-well plate method described in "Validation of two-material combinations". The bagged, stabilized powders were exposed to 50 °C for 24 hours in a static incubator and then kept in the dark at 4 °C until used.

Nodulation of *Medicago truncatula* A17 was assayed according to a protocol from Sadowsky et al. modified to growth in pouches[3–5]. Specifically, *M. truncatula* A17 seeds (Noble Research Institute) were scarified by submerging in concentrated sulfuric acid for 8-9 minutes, washed 8X with sterile water, sterilized by submerging in 8.25% sodium hypochlorite for 90 seconds, washed 8X with sterile water, allowed to imbibe in sterile water for ~1 hour, placed on 1% agar plates, sealed with parafilm and foil, vernalized at 4 °C in the dark for 5 days and germinated at 23 °C in the dark for 20 hours. Germinated seeds with root lengths of ~1.5 cm were transferred to sterilized growth pouches (Mega International, CYG) pre-wetted with 5 mL of 1/2X BNM medium (see Table S3). Five seeds were placed in each pouch. Pouches were placed between 1 inch foam blocks in a domed seed propagator (EarlyGrow, B07HHR5DGN) and incubated in plant growth chamber (Fisher Scientific, PR505755L) set to a day/night cycle of 25 °C for 16 hours and 21 °C for 8 hours. The chamber was kept humid with a large tray of water.

Two days after the seeds were transferred to the pouches, the resulting seedlings were each inoculated with 1 mL of the bacterial samples in 1/2X BMN. Specifically, the heat-stressed stabilized *E. meliloti* powders were rehydrated in 1/2X BNN at a common concentration equivalent to $10^8$ CFU/mL of the initial bacterial culture before freeze drying. Similarly, freshly grown *E. meliloti* cells were pelleted and resuspended in 1/2X BNM to a concentration of $10^7$ CFU/mL.

The pouches were left undisturbed for 6 days, watered with 5 mL of 1/2X BNM as necessary and then inspected for nodules every other day. On day 12, the number of nodulated seedlings in each pouch was counted. The percent nodulated seedlings was calculated by dividing the number of nodulated seedlings by the total number of seedlings in the pouch.

Scanning electron microscopy of dry stabilized microbial materials (Figure 4B)

The various dry microbial materials were imaged in high vacuum mode using the secondary electron detector from the Hitachi FlexSEM TM-1000 II (Tokyo, Japan). Low voltage imaging (3 kV) was used to prevent damage from electron bombardment while providing high surface detail. The powders were mounted using double sided carbon tape (Ted Pella Inc). Using JEOL USA's Smart Coater (Peabody, MA USA), gold was coated for approximately 2.5 minutes (~4 nm of gold deposited). This conductive coating prevented excessive surface charging artifacts in images. Typical imaging conditions included a spot intensity of 50 (based on the instruments unitless scale from 1-100) and a working distance of less than 7 mm.

Data analysis

Numerical data was analyzed and plotted using Prism 9 (GraphPad). Statistical tests, replicate numbers and errors are indicated for each relevant figure panel.

**A**

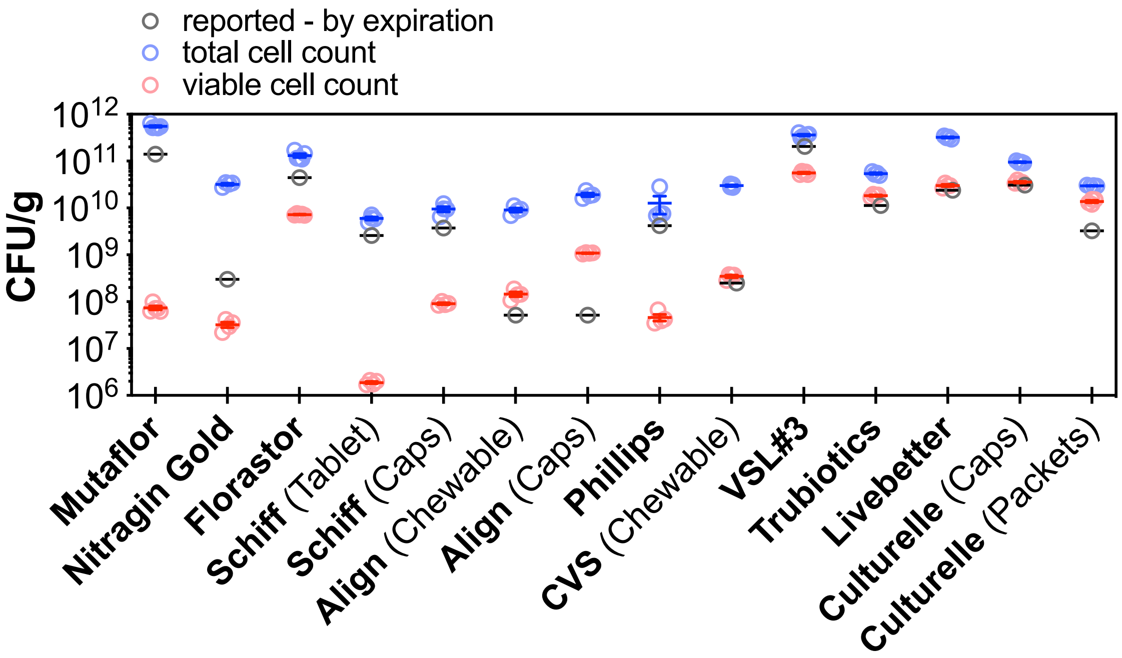

**B**

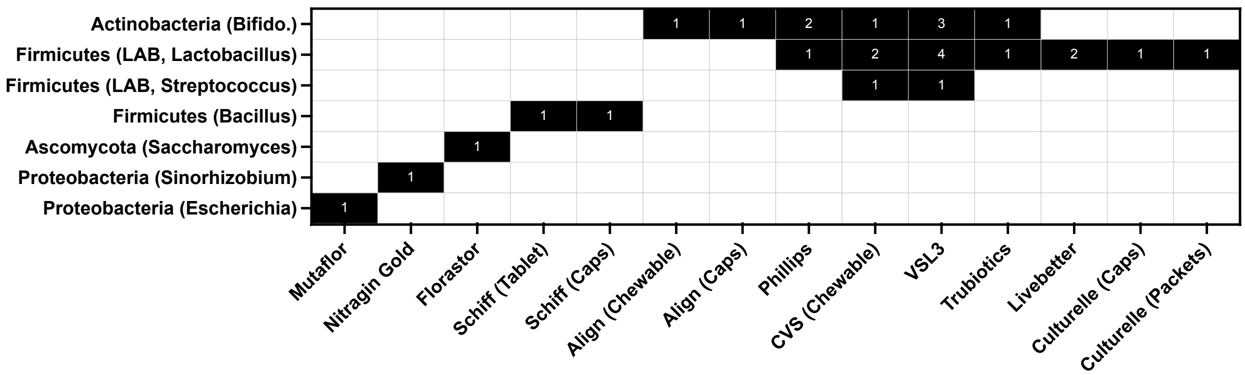

**Fig. S1. Raw CFU/g values of probiotics survey. A.** Raw data used to calculate the percent viabilities graphed in Figure 1. Means and standard errors of the mean (SEM) are plotted over individual replicates. **B.** Phylogenetic composition of the commercial products assessed. Numbers in boxes indicate the number of distinct strains in each clade. Full product details are summarized in Table S1.

**A**

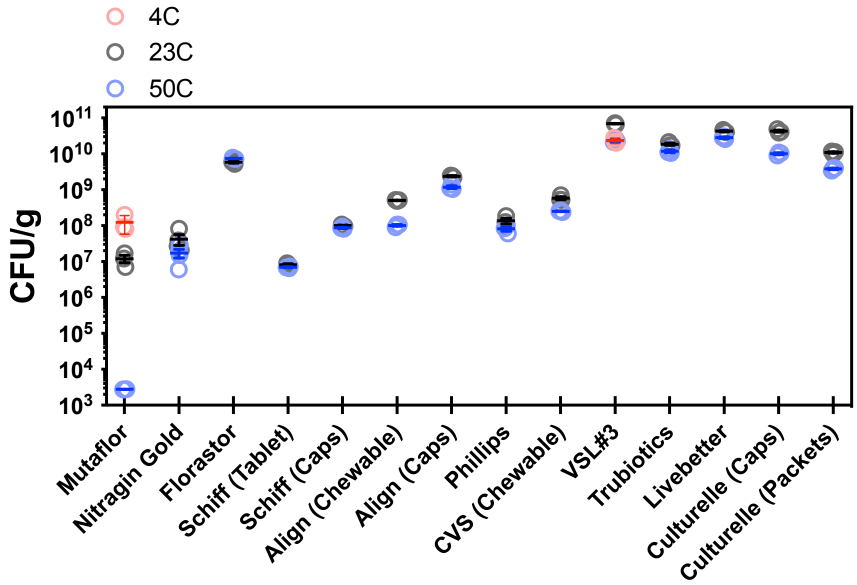

**B**

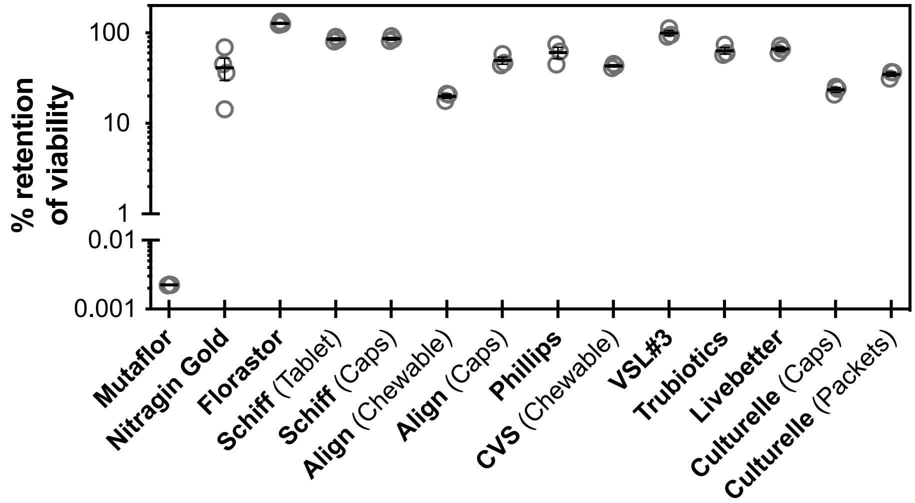

**Fig. S2. Commercial probiotic product stress tested at 50 C for 24 hours.** A. Each product was kept at 4 °C, 23 °C or 50 °C for 24 hours and the resulting viability was assessed (see Methods). Only Mutaflor and VSL#3 were kept at 4 °C per manufacturer recommendations. B. Analysis of data in panel A. All viability values at 50 °C were divided by the mean viability at the manufacturer recommended storage temperature (23 °C for all except for Mutaflor and VSL#3 were the value from the 4 °C samples were used). Means and standard errors of the mean (SEM) are plotted over individual replicates for both panels.

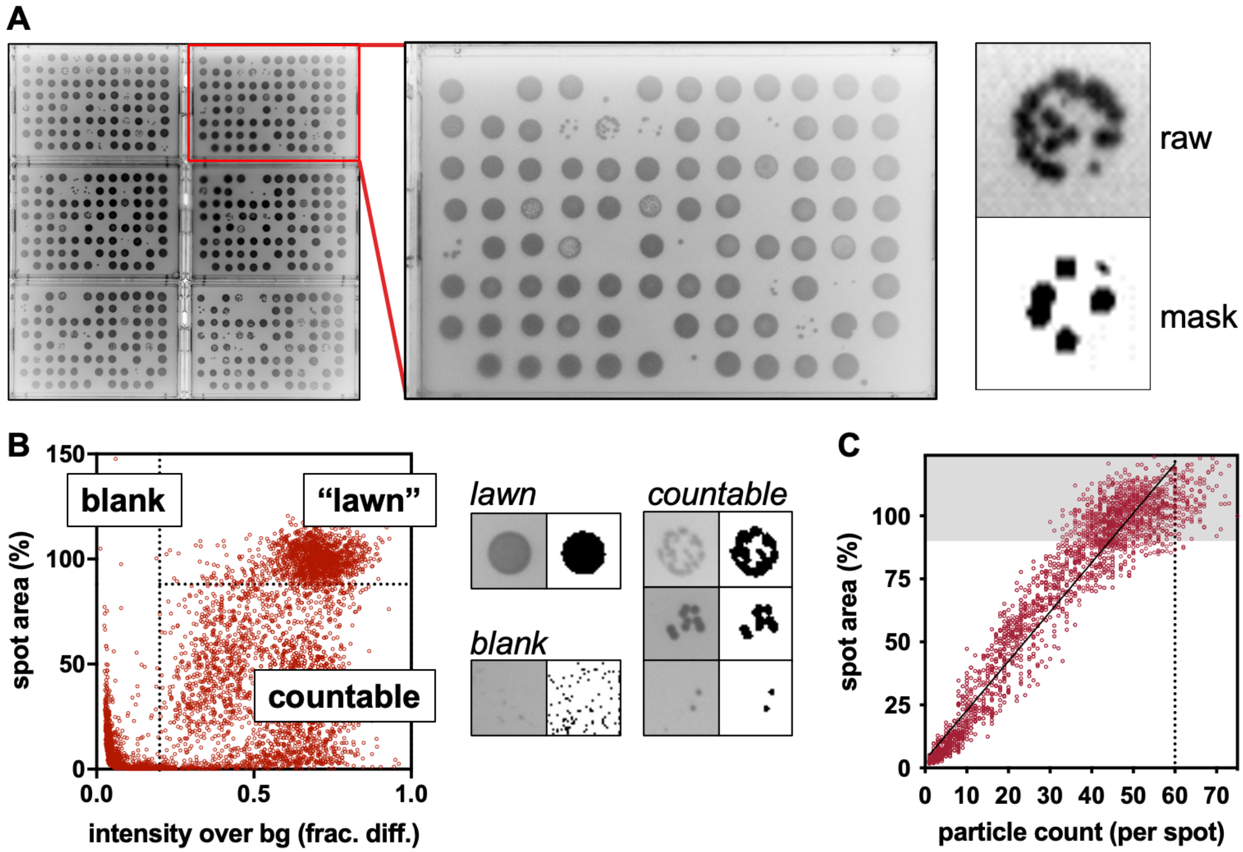

**Fig. S3. High throughput pipeline for assessing viability score. A.** Dried bacterial samples are rehydrated, diluted, plated onto 1-well plates in batch and incubated. The resulting array of spots is imaged, and each spot image ("raw") is programmatically segmented ("mask") into a region of interest (i.e., region containing bacterial colonies) and the segmented region is subdivided into countable "particles" using a watershed algorithm using Fiji. **B.** Additional image parameters from the segmented region such as the size relative to a control lawn size (spot are percent) and mean signal intensity relative to the region outside the segmented region (intensity over background) can be used to automatically classify spots into "lawns", "blanks" and "countable" spots. The particle counts for blank spots are set to zero. The particle counts of countable spots are interpreted as the number of colonies. **C.** The particle counts of lawns are set to an organism specific value determined by extrapolating the line of best fit of all countable spots (i.e. those below grey region) to a particle count value corresponding to a spot area percent of 120%.

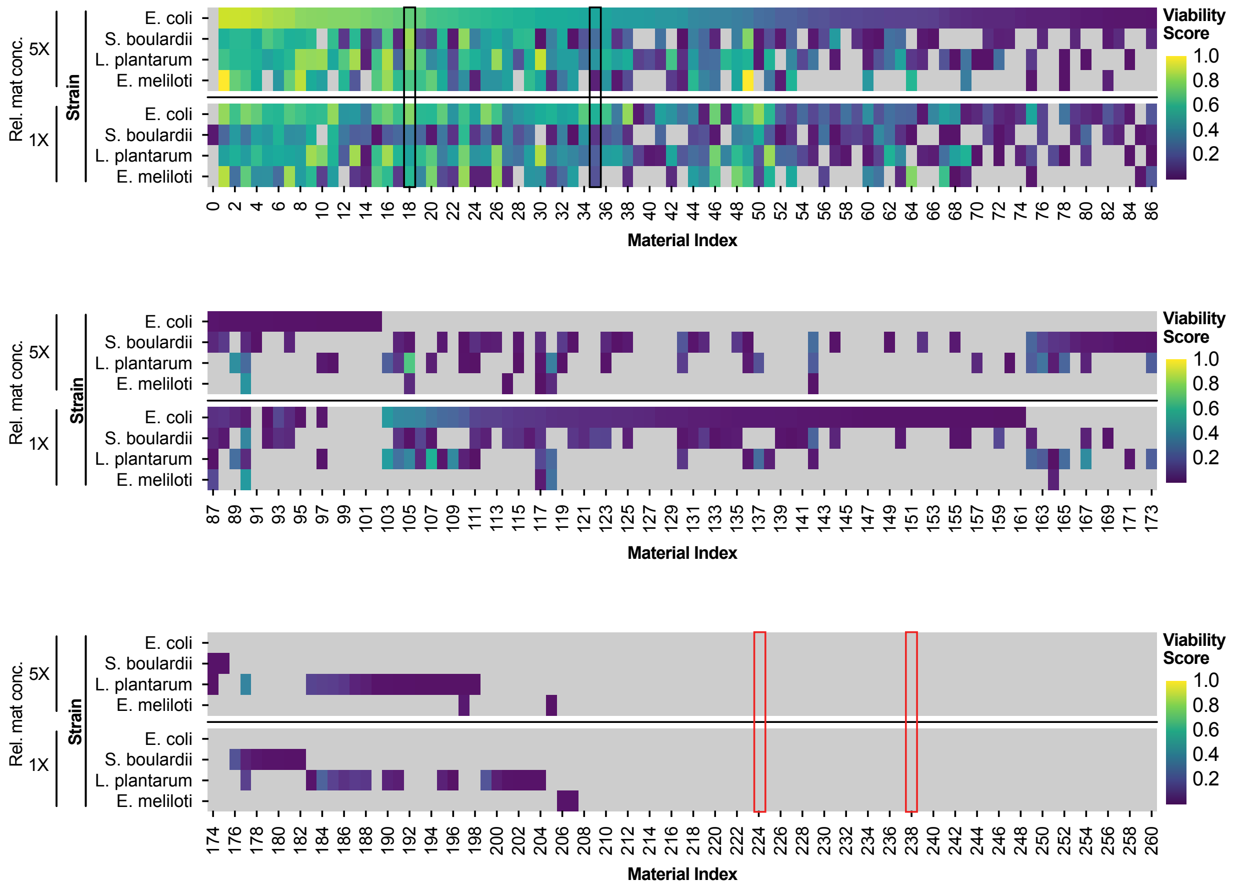

**Fig. S4. Detailed results of material stabilizer library.** Extended data of Figure 2C plotted with a continuous color scale. Grey color denotes zero observed viability. Vertical black boxes mark the positive controls (#18: ATCC reagent 20 and #35: trehalose) and vertical red boxes mark the negative controls (#224: sodium hydroxide and #238: sodium metabisulfite). Viability score is a composite score of the colony counts across three plating dilutions (see Methods). "Rel. mat. conc." – relative material concentrations as defined in Table S2 for 5X and 1X.

x

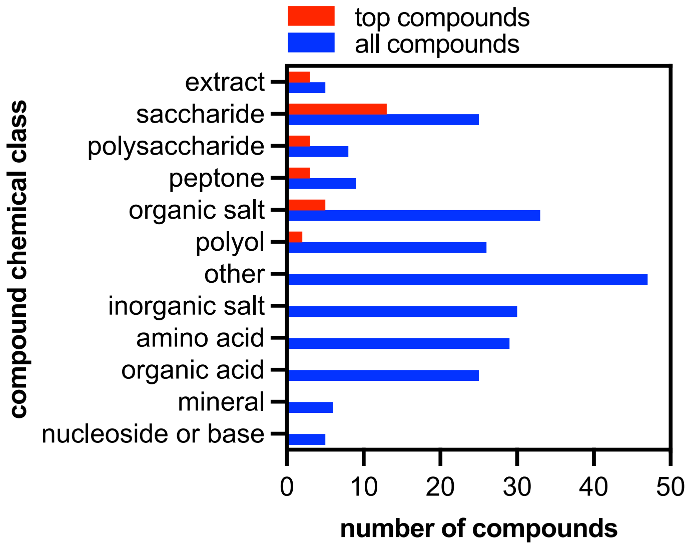

**Fig. S5. Compound classes over-represented in hit materials.** All compounds were classified by chemical structure. Top performing compounds are the same ones included in the correlation analysis (Fig. 2D, see Methods). Extract refers to cell or plant extracts such as yeast extract or malt extract. Peptone refers to both protein digests (tryptone) or other protein-based materials (catalase). Mixtures of individually prepared components were omitted from this analysis.

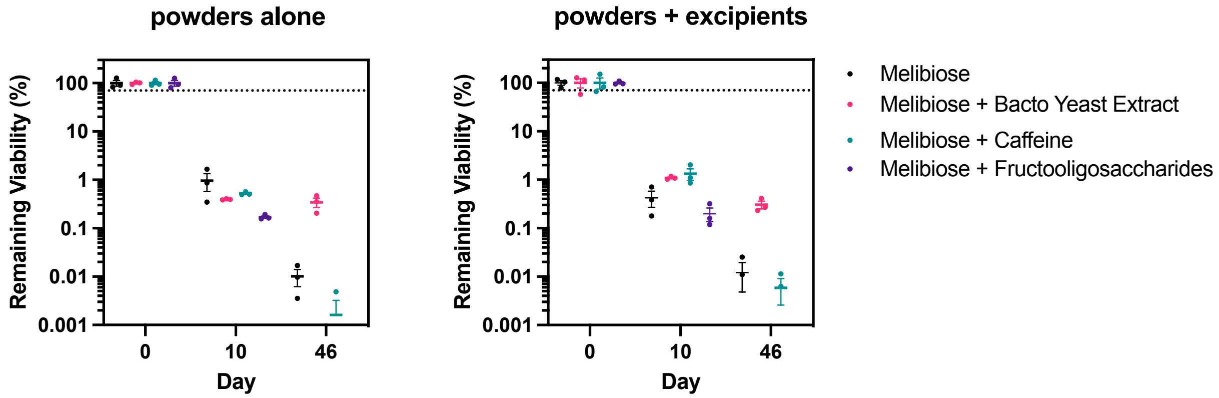

**Fig. S6. Validation of two-material formulations at 37 °C.** Materials selected from the two-material library were validated at a larger scale (see Methods) to determine the precise retention of viability when stored at 37 °C for the specified time. "Powders alone" refers to storage of the as freeze-dried and milled powders in nitrogen-flushed bags with desiccant. "Powders + excipients" refers to storage of the milled microbial powders mixed with excipients (binder, filler, glidant) as described in the Methods). Means and standard errors of the mean (SEM) are plotted over individual replicates for both panels.

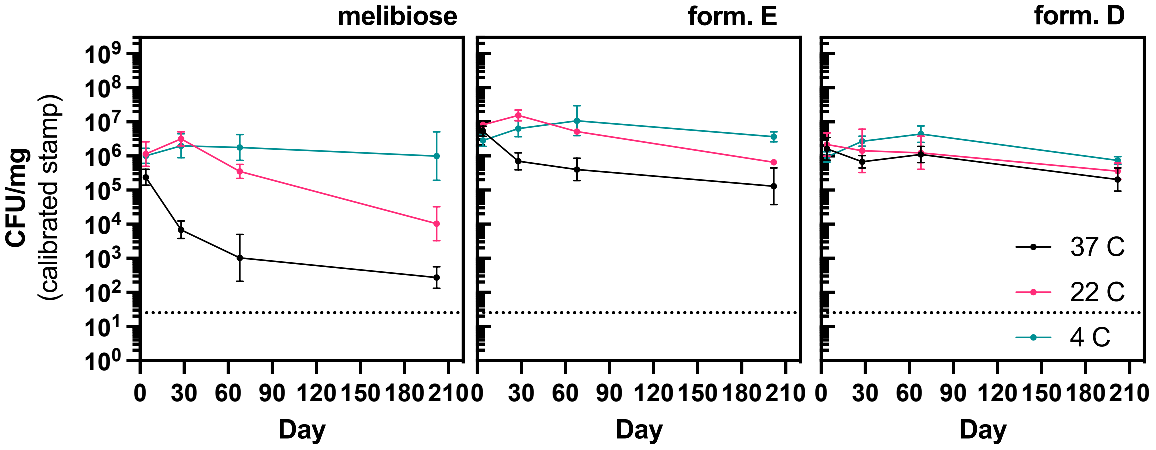

**Fig. S7. Individual viability traces of top materials formulations at different storage temperatures.** Extended data from Figure 2D. Freeze-dried microbial powders were stored after milling and mixing with excipients (see Methods). Storage was in glass vials with desiccant in the dark at the specified temperatures. Lines connect the geometric means and error bars mark the 95% confidence intervals. N = 3.

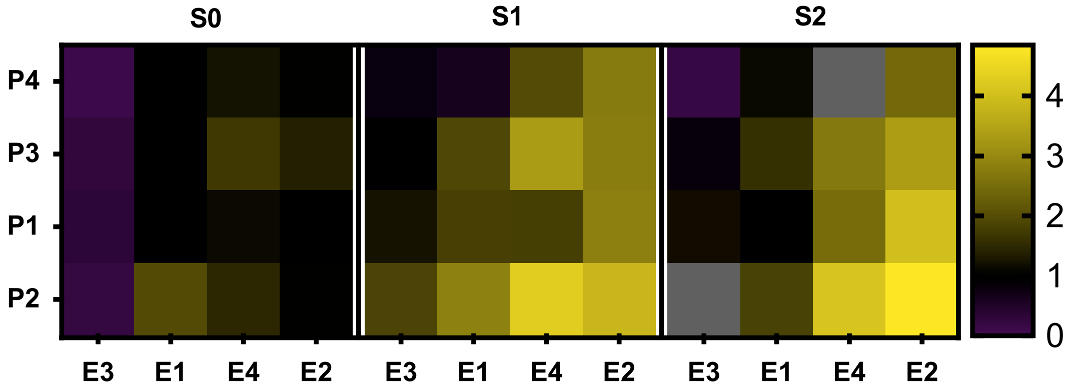

**Fig. S8. Preculture medium impacts ultimate survival through lyophilization.** *E. coli* Nissle 1917 was grown in a range of preculture media, washed in PBS, mixed with trehalose, freeze-dried, hydrated, inoculated in fresh LB and then mixed with Presto Blue to assess the viability. Values represent the fold increase in Presto Blue signal relative to the control medium (LB). Each medium has three components a Peptone (P#) at 10 g/L, an Extract (E#) at 5 g/L and a Salt (NaCl) concentration (S#). P1: Tryptone, P2: Soytone, P3: Gelysate peptone, P4: Bacto peptone, E1: Beef extract, E2: Yeast extract, E3: Malt extract, E4: Beef & yeast extract, S0: 86 mM NaCl, S1: 250 mM NaCl, S2: 600 mM NaCl. The control LB well is P1:E2:S0.

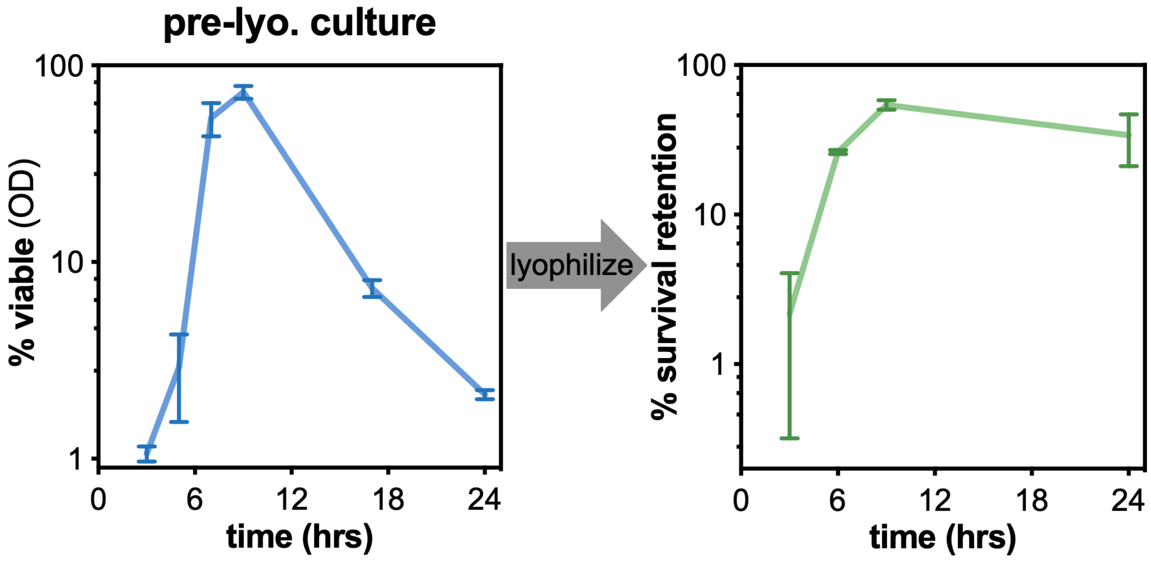

**Fig. S9. Time of bacterial harvest impacts ultimate survival through lyophilization.** Liquid flask cultures of *E. coli* Nissle 1917 were harvested at the specified times since inoculation. The viability of the culture at the time of harvest was assessed by plating and a percent viability was calculated by dividing by the estimated CFU / mL estimated from the optical density measurement (1 $OD_{600}$ ~ 1E9 CFU / mL). Each sample of harvested cells was freeze-dried and the resulting viability was assessed after 24 hours of storage at 23 °C (see Methods). Lines connect the means and error bars mark the standard error of the means. N = 2.

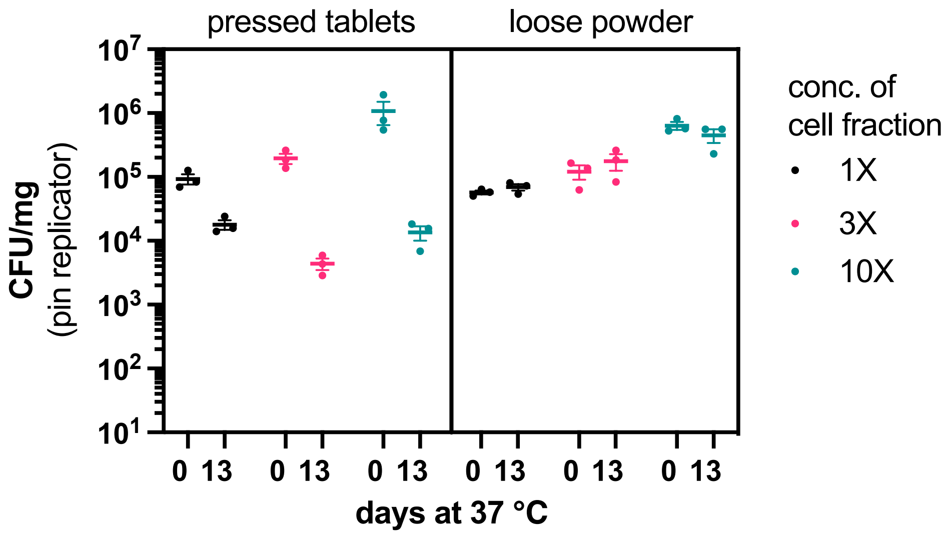

**Fig. S10. Bacterial loading can be increased in powders and tablets.** *E. coli* Nissle 1917 was freeze dried with Formulation D at three different concentrations of the initial cell suspension. The resulting powders were milled and mixed with excipients (see Methods) and stored as mixed (right) or tableted and then stored (left). All samples were stored in nitrogen-flushed bags with desiccant at 37 °C for the specified time. Means and standard errors of the mean (SEM) are plotted over individual replicates.

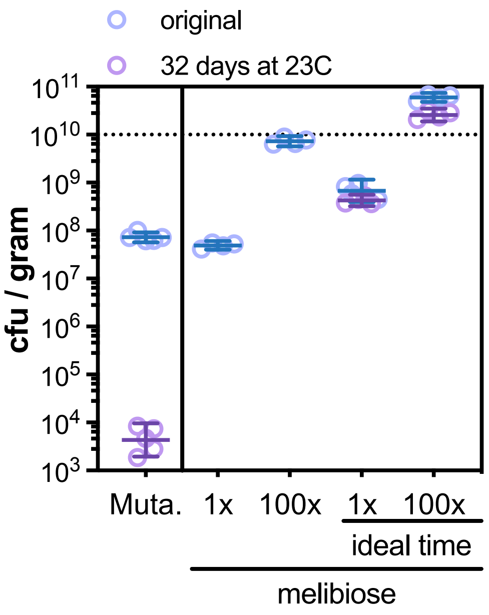

**Fig. S11. Combination of preculture timing and increased bacterial loading leads to viabilities above $10^{10}$ cfu per gram.** Either the commercial product Mutaflor or the synthetic extremophile *E. coli* Nissle 1917 (melibiose 1X) were evaluated for their initial viability and viability after long-term storage at 23 °C for 32 days. "100X" refers to a 100-fold increase in the bacterial concentration of the bacterial suspension mixed with the stabilizer. "ideal time" refers to harvesting the bacterial flask culture at the optimal time of 9 hours as indicated by the data in Figure S10. Geometric means and 95% confidence intervals are plotted over individual replicates.

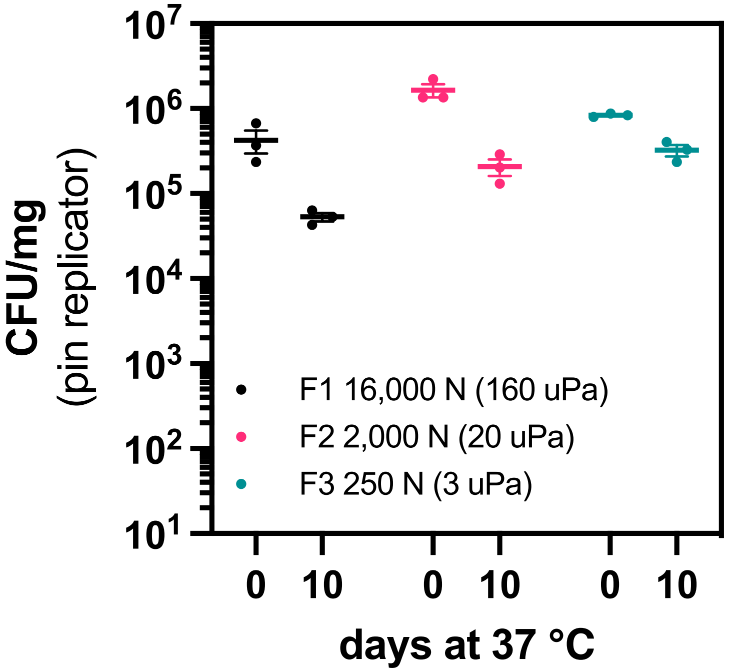

**Fig. S12. Tableting pressure modulates stability of bacteria at 37 °C.** *E. coli* Nissle 1917 was freeze dried with Form. D at a 10X cell concentration of the initial bacterial suspension. The resulting material was milled, mixed with excipients, and pressed into tablets (see Methods) at the pressure indicated). Means and standard errors of the mean (SEM) are plotted over individual replicates.

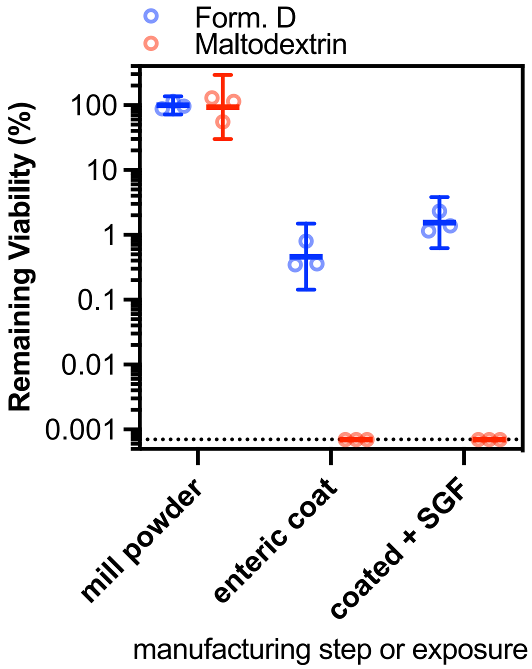

**Fig. S13. Synthetic extremophile E. coli Nissle 1917 extremophile survives in SGF for 1 hour.** Extended data from Figure 4C. Synthetic extremophile *E. coli* Nissle 1917 (Form D) or the commercial comparator (freeze-dried with 5% maltodextrin) were milled, tableted and coated with Eudragit S100 (see Methods). The coated tablets were subsequently submerged in simulated gastric fluid (SGF) for 1 hour at 37 °C to simulate ingestion and passage through the stomach. The viability of each sample was assessed by plating. Viability is normalized to the viability of the milled powder. Coated tablets were cut in half immediately before rehydration to expose inner contents. Geometric means and 95% confidence intervals are plotted over individual replicates. Horizontal dotted line marks the limit of detection.

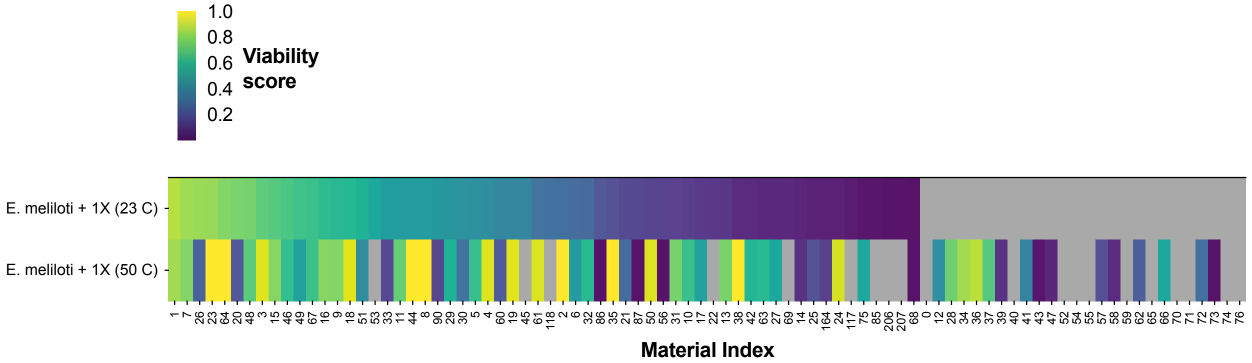

**Fig. S14.** ***E. meliloti*** **stabilizers against 24 hours at 50 °C.** Extended data of Figure S4. *E. meliloti* was mixed with the material library at the 1X concentration, freeze dried, and exposed to 50 °C for 24 hours. The viability score was characterized as in Figure S4, and the data is ordered by the viability score of the comparison values when stored at 23 °C.

### Table S1. Commercial probiotic and microbial materials analyzed

*"Days until exp." is the number of days until the expiration date on the day the assessment was carried out. "Mass per dose" is the mean mass of the microbial fraction of the doses used in the assessment (see Methods). "Hyd. vol. per dose" is the volume used to rehydrate each dose. All culture media solidified with 1.5% agar (details are listed in Table S3).*

| Product Name | Lot # | Exp. date | Days until exp. | Mass per dose | Hyd. vol. per dose | Culture conditions (medium, temp, $O_2$) |
|---|---|---|---|---|---|---|
| *Details for Figures 1, S1* | | | | | | |
| Mutaflor | 830210 | 8/15/19 | 161 | 0.18 g | 50 mL | LB, 37℃, aerobic |
| Nitragin Gold Alfalfa and Sweet Clover | NGA19123 | 1/31/22 | 903 | 0.05 g | 4 mL | TY, 30℃, aerobic |
| Florastor | 2033 | 5/1/21 | 816 | 0.28 g | 50 mL | YPD, 30℃, aerobic |
| Schiff Digestive Advantage Prebiotic Fiber Plus Probiotic Tablets | 4345700 | 4/1/20 | 266 | 0.31 g | 10 mL | Nutrient, 37℃, aerobic |
| Schiff Digestive Advantage Daily Probiotic Capsules | 4167300 | 5/1/20 | 305 | 0.54 g | 50 mL | Nutrient, 37℃, aerobic |
| Align Probiotic Chewables for Adults Supplement | 83121453A2 | 8/1/20 | 396 | 0.19 g | 25 mL | MRS+cys, 37℃, anaerobic |
| Align Probiotic Supplement 24/7 Digestive Support | 83471453A1 | 9/1/20 | 448 | 0.19 g | 25 mL | MRS+cys, 37℃, anaerobic |
| Phillips' Colon Health Probiotic Capsules | 8H17A | 8/1/20 | 400 | 0.36 g | 25 mL | MRS+cys, 37℃, anaerobic |
| CVS Health Children's Chewable Probiotic Tablets | TA244 | 2/1/21 | 545 | 0.32 g | 50 mL | MRS, 37℃, anaerobic |
| VSL#3 | 806084 | 6/1/20 | 441 | 0.55 g | 50 mL | MRS, 37℃, anaerobic |
| TruBiotics Capsules | 55143 | 5/1/20 | 318 | 0.18 g | 50 mL | MRS, 37℃, anaerobic |
| Live Better Adult Advanced Daily Probiotics | SA004 | 2/1/20 | 227 | 0.5 g | 50 mL | MRS, 37℃, anaerobic |
| Culturelle Digestive Daily Probiotic Capsules | 19003CLG2 | 12/1/20 | 538 | 0.33 g | 50 mL | MRS, 37℃, anaerobic |
| Culturelle Kids Purely Probiotics Packets | 19051C3M11 | 11/1/21 | 865 | 1.5 g | 50 mL | MRS, 37℃, anaerobic |
| *Details for Figures S2 (high temperature stress test)* | | | | | | |
| Mutaflor | 910230 | 2/4/20 | 197 | 0.18 g | 50 mL | LB, 37℃, aerobic |
| Nitragin Gold Alfalfa and Sweet Clover | NGA19123 | 1/31/22 | 929 | 0.05 g | 4 mL | TY, 30℃, aerobic |
| Florastor | 2049 | 8/1/21 | 741 | 0.29 g | 50 mL | YPD, 30℃, aerobic |
| Schiff Digestive Advantage Prebiotic Fiber Plus Probiotic Tablets | 4345700 | 4/1/20 | 240 | 0.38 g | 50 mL | Nutrient, 37℃, aerobic |
| Schiff Digestive Advantage Daily Probiotic Capsules | 4167300 | 5/1/20 | 269 | 0.53 g | 50 mL | Nutrient, 37℃, aerobic |
| Align Probiotic Chewables for Adults Supplement | 83121453A2 | 8/1/20 | 368 | 0.19 g | 50 mL | MRS+cys, 37℃, anaerobic |
| Align Probiotic Supplement 24/7 Digestive Support | 83471453A1 | 9/1/20 | 419 | 0.19 g | 50 mL | MRS+cys, 37℃, anaerobic |
| Phillips' Colon Health Probiotic Capsules | 8H17A | 8/1/20 | 359 | 0.35 g | 50 mL | MRS+cys, 37℃, anaerobic |
| CVS Health Children's Chewable Probiotic Tablets | TA244 | 2/1/21 | 545 | 0.32 g | 50 mL | MRS, 37℃, anaerobic |
| VSL#3 | 806084 | 6/1/20 | 312 | 0.54 g | 50 mL | MRS, 37℃, anaerobic |
| TruBiotics Capsules | 55143 | 5/1/20 | 296 | 0.18 g | 50 mL | MRS, 37℃, anaerobic |
| Live Better Adult Advanced Daily Probiotics | SA004 | 2/1/20 | 198 | 0.5 g | 50 mL | MRS, 37℃, anaerobic |
| Culturelle Digestive Daily Probiotic Capsules | 19003CLG2 | 12/1/20 | 502 | 0.32 g | 50 mL | MRS, 37℃, anaerobic |
| Culturelle Kids Purely Probiotics Packets | 19051C3M11 | 11/1/21 | 830 | 1.5 g | 50 mL | MRS, 37℃, anaerobic |

**Table S2. Library of materials used in stabilizer screen**

*"Solution" notes when materials were purchased or procured as defined solutions and used as is. 1X concentrations were made by volumetrically diluting 5X solutions 1:4 in ultrapure water. Recipe for ATCC reagents at end of table.*

| Index | Material name | CAS or product # | Vendor | Product # | 5X conc. (% w/w) |
|---|---|---|---|---|---|
| 1 | Bacto soytone | BD 243620 | BD | 243620 | 29 |
| 2 | ATCC Reagent 18 | ATCC recipe | - | - | solution |
| 3 | Palatinose hydrate | 343336-76-5 | Sigma | P2007 | 32 |
| 4 | Difco YPD Broth | BD 242820 | BD | 242820 | 29 |
| 5 | D-(+)-Turanose | 547-25-1 | Sigma | T2754 | 40 |
| 6 | maltitol | 585-88-6 | Sigma | M8892 | 44 |
| 7 | Difco Lactobacilli MRS Broth | BD 288130 | BD | 288130 | 29 |
| 8 | OPS MFDB | OPS 500-06 | OPS | 500-06 | solution |
| 9 | sucrose | 57-50-1 | Sigma | S7903 | 50 |
| 10 | Bacto beef extract, dessicated | BD 211520 | BD | 211520 | 29 |
| 11 | Melibiose | 585-99-9 | Sigma | M5500 | 44 |
| 12 | L-Rhamnose monohydrate | 10030-85-0 | Sigma | 83650 | 29 |
| 13 | Bacto tryptone | BD 211705 | BD | 211705 | 29 |
| 14 | D-(+)-Galactose | 59-23-4 | Sigma | G6404 | 40 |
| 15 | D-(+)-Melezitose monohydrate | 10030-67-8 | Sigma | 63620 | 25 |
| 16 | Bacto malt extract | BD 218630 | BD | 218630 | 39 |
| 17 | D(-)-Fructose | 57-48-7 | Acros Organics | 161350010 | 24 |
| 18 | ATCC Reagent 20 | ATCC recipe | - | - | solution |
| 19 | 1-Kestose | 470-69-9 | Sigma | 72555 | 7.4 |
| 20 | Maltodextrin 16.5-19.5 DE | 9050-36-6 | Sigma | 419699 | 44 |
| 21 | Glucose | 50-99-7 | Sigma | G8270 | 42 |
| 22 | 2-Deoxy-D-glucose | 154-17-6 | Sigma | D3179 | 23 |
| 23 | L-glutamic acid monosodium salt monohydrate | 6106-04-3 | Sigma | 49621 | 44 |
| 24 | Maltose monohydrate | 6363-53-7 | Sigma | M5885 | 3.9 |
| 25 | (+)-Sodium L-ascorbate | 134-03-2 | Sigma | A7631 | 21 |
| 26 | polydextrose | Honeyville 77-121 | Honeyville | 77-121 | 50 |
| 27 | D-(+)-Mannose | 3458-28-4 | Sigma | M6020 | 17 |
| 28 | D-(+)-Raffinose pentahydrate | 17629-30-0 | Sigma | 83400 | 40 |
| 29 | lactulose | 4618-18-2 | Sigma | 61360 | 38 |
| 30 | Gelysate peptone | BD 211870 | BD | 211870 | 29 |
| 31 | D-cellobiose | 528-50-7 | Sigma | C7252 | 6.3 |
| 32 | Sodium gluconate | 527-07-1 | Sigma | S2054 | 32 |
| 33 | Barium acetate | 543-80-6 | Sigma | 243671 | 32 |
| 34 | sodium citrate, dihydrate | 6132-04-3 | Mallinckrodt | 0754 | 33 |
| 35 | Trehalose dihydrate | 6138-23-4 | Sigma | T9531 | 3.9 |
| 36 | alpha-lactose, monohydrate | 5989-81-1 | Sigma | L2643 | 14 |
| 37 | Difco LB Broth, Lennox | BD 240230 | BD | 240230 | 29 |
| 38 | Difco M17 Broth | BD 218561 | BD | 218561 | 29 |
| 39 | Maltodextrin 4-7 DE | 9050-36-6 | Sigma | 419672 | 44 |
| 40 | 2-Phospho-L-ascorbic acid trisodium salt | 66170-10-3 | Sigma | 49752 | 33 |
| 41 | Difco Middlebrook 7H9 Broth | BD 271310 | BD | 271310 | 29 |
| 42 | chondroitin sulfate A | 39455-18-0 | Sigma | C8529 | 7.4 |
| 43 | Karaya Gum | 9000-36-6 | Sigma | G0503 | 1 |
| 44 | Bacto yeast extract | BD 212750 | BD | 212750 | 33 |
| 45 | Xylitol | 87-99-0 | Sigma | X3375 | 34 |
| 46 | beta-lactose | 5965-66-2 | Sigma | L3750 | 44 |
| 47 | Casamino acids | BD 228820 | BD | 228820 | 1.6 |
| 48 | Bacto Tryptic Soy Broth | BD 211825 | BD | 211825 | 29 |
| 49 | Skim milk powder | 999999-99-4 | sigma | 1153630500 | 29 |
| 50 | Potassium gluconate | 299-27-4 | Sigma | P1847 | 33 |
| 51 | D-Sorbitol | 50-70-4 | Sigma | S6021 | 55 |

| | | | | | |
|---|---|---|---|---|---|
| 52 | L-(+)-Arabinose | 5328-37-0 | Sigma | A3256 | 44 |
| 53 | Adonitol | 488-81-3 | Sigma | A5502 | 34 |
| 54 | Magnesium phosphate dibasic trihydrate | 7782-75-4 | Sigma | 63080 | 30 |
| 55 | Thymidine | 50-89-5 | Sigma | T9250 | 5.6 |
| 56 | sodium phosphate, monobasic | 7558-80-7 | Sigma | S8282 | 40 |
| 57 | γ-Aminobutyric acid | 56-12-2 | Sigma | A2129 | 50 |
| 58 | myo-Inositol | 87-89-8 | Sigma | I7508 | 9.9 |
| 59 | L-glutamic acid | 56-86-0 | Sigma | G8415 | 0.69 |
| 60 | Calcium D-gluconate | 299-28-5 | Sigma | C8231 | 2.5 |
| 61 | Potassium citrate tribasic monohydrate | 6100-05-6 | Sigma | C3029 | 55 |
| 62 | Lithium acetate dihydrate | 6108-17-4 | Sigma | L4158 | 40 |
| 63 | Cytidine | 65-46-3 | Sigma | C122106 | 7.4 |
| 64 | Difco Nutrient Broth | BD 234000 | BD | 234000 | 29 |
| 65 | Poly(vinylpolypyrrolidone) | 9003-39-8 | Sigma | 77627 | 7.4 |
| 66 | L-Histidine | 71-00-1 | Sigma | H8000 | 3.3 |
| 67 | Magnesium D-gluconate hydrate | 3632-91-5 | Sigma | M7554 | 9.1 |
| 68 | Aluminum silicate | Sigma 520179 | Sigma | 520179 | 40 |
| 69 | Lecithin, Refined | 8002-43-5 | Alfa Aesar | 36486 | 15 |
| 70 | Celite® 545 | Sigma CX0574-1 | Sigma | CX0574-1 | 11 |
| 71 | 1,4-Piperazinediethanesulfonic acid | 5625-37-6 | Sigma | P1851 | 10 |
| 72 | Amygdalin | 29883-15-6 | Sigma | 10050 | 7.4 |
| 73 | Catalase from bovine liver | 9001-05-2 | Sigma | C9322 | 0.07 |
| 74 | Ammonium sulfate | 7783-20-2 | Mallinckrodt | 3512 | 38 |
| 75 | OPS Lyophilization reagent | OPS 500-02 | OPS | 500-02 | solution |
| 76 | α-Cyclodextrin | 10016-20-3 | Sigma | C4642 | 0.79 |
| 77 | Lithium chloride | 7447-41-8 | Sigma | 62478 | 35 |
| 78 | D-glucosamine HCl | 66-84-2 | Sigma | G1514 | 7.4 |
| 79 | Catalase (Aspergillus niger) | 9001-05-2 | Sigma | C3515 | 0.07 |
| 80 | N,N-Bis(2-hydroxyethyl)-2-aminoethanesulfonic acid | 10191-18-1 | Sigma | B9879 | 50 |
| 81 | Yeast nitrogen base | MP 4027512 | MP biomedicals | 4027512 | 9.1 |
| 82 | Adenosine | 58-61-7 | Sigma | A9251 | 0.53 |
| 83 | Hypoxanthine | 68-94-0 | Sigma | H9636 | 0.06 |
| 84 | 1-Adamantylamine | 768-94-5 | Sigma | 138576 | 0.5 |
| 85 | Mucin (Porcine, type III) | 84082-64-4 | Sigma | M1778 | 0.4 |
| 86 | Allantoin | 97-59-6 | Sigma | 05670 | 0.42 |
| 87 | 3,4-dihydroxy-DL-phenylalanine | 63-84-3 | Sigma | D9503 | 0.4 |
| 88 | hydrocortisone | 50-23-7 | Sigma | H4001 | 0.02 |
| 89 | Hydroxyapatite nanopowde | 12167-74-7 | Sigma | 677418 | 23 |
| 90 | Sodium caseinate | 9005-46-3 | Sigma | C8654 | 3.9 |
| 91 | O-Phospho-DL-serine | 17885-08-4 | Sigma | 79710 | 1.5 |
| 92 | N-Phenylthiourea | 103-85-5 | Sigma | P7629 | 0.2 |
| 93 | N-Hydroxyphthalimide | 524-38-9 | Sigma | H53704 | 0.25 |
| 94 | L-Tyrosine | 60-18-4 | Sigma | T3754 | 0.03 |
| 95 | 1-Pentanol | 71-41-0 | Sigma | 138975 | 2.2 |
| 96 | sodium acetate, anhydrous | 127-09-3 | Mallinckrodt | 7372 | 27 |
| 97 | Ethylenediaminetetraacetic acid | 60-00-4 | Sigma | 03690 | solution |
| 98 | 2,2-Bis(hydroxymethyl)propionic acid | 4767-03-7 | Sigma | 106615 | 9.1 |
| 99 | Pimelic acid | 111-16-0 | Sigma | P45001 | 3.9 |
| 100 | 2,4-Pentanediol | 625-69-4 | Sigma | 156019 | 50 |
| 101 | 1-Thioglycerol | 96-27-5 | Sigma | M6145 | 50 |
| 102 | 1,5,7-Triazabicyclo[4.4.0]dec-5-ene | 5807-14-7 | Sigma | 345571 | 9.1 |
| 103 | L-Arginine | 74-79-3 | Sigma | A5006 | 13 |
| 104 | Choline bitartrate | 87-67-2 | Sigma | C1629 | 7.4 |
| 105 | Bacto peptone | BD 211677 | BD | 211677 | 29 |
| 106 | L-serine | 56-45-1 | Sigma | S4500 | 3.9 |
| 107 | Meglumine | 6284-40-8 | Sigma | M9179 | 44 |
| 108 | L-Threonine | 72-19-5 | Sigma | T8625 | 7.2 |
| 109 | Tris(hydroxymethyl)aminomethane | 77-86-1 | Sigma | T6791 | 31 |
| 110 | erythritol | 149-32-6 | Sigma | E7500 | 33 |

| | | | | | |
|---|---|---|---|---|---|
| 111 | 2-Ethyl-1-hexanol | 104-76-7 | Sigma | 04050 | 0.07 |
| 112 | uracil | 66-22-8 | MP biomedicals | 4061212 | 0.29 |
| 113 | caffeine | 58-08-2 | Sigma | C0750 | 1.7 |
| 114 | ricinoleic acid | 141-22-0 | Sigma | 83903 | 0.28 |
| 115 | Suberic acid | 505-48-6 | Sigma | S5200 | 0.13 |
| 116 | Neohesperidin dihydrochalcone | 20702-77-6 | Sigma | N8757 | 0.08 |
| 117 | D-(+)-Xylose | 58-86-6 | Sigma | 95729 | 50 |
| 118 | Calcium acetate hydrate | 114460-21-8 | Sigma | C1000 | 14 |
| 119 | 6-Aminocaproic acid | 60-32-2 | Sigma | A2504 | 29 |
| 120 | Sucrose octaacetate | 126-14-7 | Sigma | W303801 | 0.06 |
| 121 | α-Cyano-4-hydroxycinnamic acid | 28166-41-8 | Sigma | 55341 | 0.48 |
| 122 | trans-cinnamic acid | 140-10-3 | Sigma | C80857 | 0.03 |
| 123 | beta-cyclodextrin | 7585-39-9 | Sigma | C4767 | 1.5 |
| 124 | N-Hydroxysuccinimide | 6066-82-6 | Sigma | 130672 | 3.9 |
| 125 | Sulfanilamide | 63-74-1 | Sigma | S9251 | 0.67 |
| 126 | boric acid | 10043-35-3 | Sigma | B6768 | 3.1 |
| 127 | Triacetin | 102-76-1 | Sigma | 525073 | 5.8 |
| 128 | Mannitol | 69-65-8 | Sigma | M4125 | 12 |
| 129 | Propyl gallate | 121-79-9 | Sigma | P3130 | 0.28 |
| 130 | Betaine | 107-43-7 | Sigma | B2629 | 57 |
| 131 | D-tyrosine | 556-02-5 | Fluka | 93840 | 0.04 |
| 132 | (−)-Terpinen-4-ol | 20126-76-5 | Sigma | 11584 | 10 |
| 133 | L-LEUCINE | 61-90-5 | MP biomedicals | 4060512 | 1.9 |
| 134 | fumaric acid | 110-17-8 | Sigma | F8509 | 0.5 |
| 135 | L-Methionine | 63-68-3 | Sigma | M9625 | 4.3 |
| 136 | Pentaerythritol | 115-77-5 | Sigma | P4755 | 5.5 |
| 137 | Zinc acetate | 557-34-6 | Sigma | 383317 | 20 |
| 138 | Tyramine | 51-67-2 | Sigma | T90344 | 0.83 |
| 139 | 4-Guanidinobutyric acid | 463-00-3 | Sigma | G6503 | 7.4 |
| 140 | Guanosine 5′-monophosphate disodium salt hydrate | 5550-12-9 | Sigma | G8377 | 3.9 |
| 141 | Taurine | 107-35-7 | Sigma | T0625 | 7.1 |
| 142 | Albumin, human | 70024-90-7 | Sigma | A1653 | 7.4 |
| 143 | β-Alanine | 107-95-9 | Sigma | 146064 | 31 |
| 144 | Saccharin | 81-07-2 | Sigma | 109185 | 0.27 |
| 145 | salicylic acid | 69-72-7 | Sigma | 247588 | 0.18 |
| 146 | 1,4-Cyclohexanedimethanol | 105-08-8 | Sigma | 125598 | 43 |
| 147 | L-alanine | 56-41-7 | sigma | A7627 | 12 |
| 148 | Ammonium bromide | 12124-97-9 | Fluka | 213349 | 39 |
| 149 | trans-4-Hydroxy-L-proline | 51-35-4 | Sigma | H54409 | 22 |
| 150 | Propyl 4-hydroxybenzoate | 94-13-3 | Sigma | P53357 | 0.03 |
| 151 | 1,4-Benzenedimethanol | 589-29-7 | Sigma | B3000 | 2.3 |
| 152 | O-tert-Butyl-L-serine | 18822-58-7 | Sigma | B6278 | 9.1 |
| 153 | Acetylsalicylic acid | 50-78-2 | Sigma | 239631 | 0.22 |
| 154 | Urea | 57-13-6 | Sigma | U4884 | 44 |
| 155 | L-Tryptophan | 73-22-3 | Sigma | T0254 | 0.9 |
| 156 | 3-Pentanol | 584-02-1 | Sigma | P8025 | 4.2 |
| 157 | 3-Guanidinopropionic acid | 353-09-3 | Sigma | G6878 | 7.4 |
| 158 | dioctyl-sulfosuccinate | 577-11-7 | Sigma | 323586 | 1.2 |
| 159 | Octanoic acid | 124-07-2 | Sigma | C2875 | 0.05 |
| 160 | Adipic acid | 124-04-9 | Sigma | 09582 | 1.8 |
| 161 | Triglycerol | 20411-31-8 | Sigma | 17782 | 33 |
| 162 | sodium triphosphate, pentabasic | 7758-29-4 | Sigma | T5883 | 14 |
| 163 | Tris(hydroxymethyl)aminomethane hydrochloride | 1185-53-1 | Sigma | T6666 | 31 |
| 164 | L-Proline | 147-85-3 | Sigma | P0380 | 57 |
| 165 | Choline chloride | 67-48-1 | Sigma | C7527 | 29 |
| 166 | L-valine | 72-18-4 | Sigma | V0500 | 4.4 |
| 167 | Potassium pyrophosphate | 7320-34-5 | Sigma | 322431 | 60 |
| 168 | Sodium dodecyl sulfate | 151-21-3 | Sigma | L3771 | 7.4 |
| 169 | β-Glycerophosphate disodium salt hydrate | 154804-51-0 | Sigma | G9422 | 3.9 |

| | | | | | |
|---|---|---|---|---|---|
| 170 | hydroquinone | 123-31-9 | Sigma | H9003 | 6.6 |
| 171 | N,N-Bis(2-hydroxyethyl)ethylenediamine | 3197-06-6 | Sigma | 480614 | 9.1 |
| 172 | 3-Methyl-1,5-pentanediol | 4457-71-0 | Sigma | 68346 | 50 |
| 173 | Sodium bicarbonate | 144-55-8 | Sigma | S6014 | 7.4 |
| 174 | sodium sulfate | 7757-82-6 | Sigma | 238597 | 18 |
| 175 | L-ascorbic acid | 50-81-7 | Sigma | A5960 | 21 |
| 176 | Aluminum L-lactate | 18917-91-4 | Sigma | 430633 | 9.1 |
| 177 | Sodium chloride | 7647-14-5 | Sigma | S7653 | 22 |
| 178 | D-(-)-Salicin | 138-52-3 | Sigma | S0625 | 3.4 |
| 179 | 1,2,4,5-Benzenetetracarboxylic acid | 89-05-4 | Sigma | B4007 | 1.1 |
| 180 | 3,6-Dimethyl-1,4-dioxane-2,5-dione | 95-96-5 | Sigma | 303143 | 0.99 |
| 181 | (±)-1,3-Butanediol | 107-88-0 | Sigma | 18940 | 50 |
| 182 | Sucralose | 56038-13-2 | Sigma | 69293 | 7.4 |
| 183 | Calcium L-lactate hydrate | 41372-22-9 | Sigma | L4388 | 14 |
| 184 | Potassium Chloride | 7447-40-7 | Mallinckrodt | 6858 | 22 |
| 185 | sodium sulfite | 7757-83-7 | Sigma | 71991 | 20 |
| 186 | sodium phosphate, dibasic | 7558-79-4 | Mallinckrodt | 7917 | 9.1 |
| 187 | Ammonium carbonate | 506-87-6 | Sigma | 207861 | 17 |
| 188 | Bentonite | 1302-78-9 | Sigma | 285234 | 1.6 |
| 189 | Taurocholic acid sodium salt hydrate | 345909-26-4 | Sigma | T4009 | 7.4 |
| 190 | Carbon, mesoporous | 1333-86-4 | Sigma | 699624 | 2.6 |
| 191 | 4,7,10-Trioxa-1,13-tridecanediamine | 4246-51-9 | Sigma | 369519 | 9.1 |
| 192 | Potassium Phosphate, monobasic | 7778-77-0 | Mallinckrodt | 7100 | 15 |
| 193 | Fructooligosaccharides | Sigma F8052 | Sigma | F8052 | 15 |
| 194 | Copper(II) sulfate | 7758-98-7 | Sigma | C1297 | 14 |
| 195 | Sodium carbonate | 497-19-8 | Sigma | 223484 | 19 |
| 196 | 3-(Diethylamino)-1,2-propanediol | 621-56-7 | Sigma | 210226 | 9.1 |
| 197 | D-(-)-Ribose | 50-69-1 | Sigma | R7500 | 40 |
| 198 | N-(1-Naphthyl)ethylenediamine dihydrochloride | 1465-25-4 | Sigma | 222488 | 0.16 |
| 199 | Potassium acetate | 127-08-2 | Sigma | 236497 | 62 |
| 200 | L-lysine | 56-87-1 | Sigma | L5501 | 44 |
| 201 | Manganese(II) chloride tetrahydrate | 13446-34-9 | Sigma | M8054 | 47 |
| 202 | DAB-Am-4 | 120239-63-6 | Sigma | 460699 | 9.1 |
| 203 | ammonium acetate | 631-61-8 | Mallinckrodt | 3272 | 44 |
| 204 | N,N′-Dimethylethylenediamine | 110-70-3 | Sigma | D157805 | 9.1 |
| 205 | citric acid, monohydrate | 5949-29-1 | Mallinckrodt | 0627 | 32 |
| 206 | 5-Aminovaleric acid | 660-88-8 | Sigma | 123188 | 44 |
| 207 | Citric acid | 77-92-9 | Sigma | 251275 | 32 |
| 208 | Guanidine hydrochloride | 50-01-1 | Sigma | G4505 | 63 |
| 209 | octyl-beta-D-glucopyranoside | 29836-26-8 | Sigma | O8001 | 7.4 |
| 210 | DL-β-(2-Thienyl)serine | 32595-59-8 | Sigma | T5000 | 0.99 |
| 211 | Cystamine dihydrochloride | 56-17-7 | Sigma | C8707 | 33 |
| 212 | mercaptosuccinic acid | 70-49-5 | Sigma | 88460 | 24 |
| 213 | Glycine | 56-40-6 | Sigma | G7126 | 17 |
| 214 | glycolic acid | 79-14-1 | Sigma | 124737 | 44 |
| 215 | Glycerol phosphate calcium salt | 58409-70-4 | Sigma | G6626 | 3.9 |
| 216 | potassium carbonate | 584-08-7 | Sigma | P5833 | 47 |
| 217 | 3-Methyl-1,3-butanediol | 2568-33-4 | Sigma | 65965 | 50 |
| 218 | cysteamine | 60-23-1 | Sigma | 30070 | 33 |
| 219 | Pyridoxine hydrochloride | 58-56-0 | Sigma | P6280 | 15 |
| 220 | L-(+)-Lactic acid | 79-33-4 | Sigma | L1750 | 7.4 |
| 221 | itaconic acid | 97-65-4 | Sigma | I29204 | 6.3 |
| 222 | Potassium hydroxide | 1310-58-3 | Sigma | P5958 | 29 |
| 223 | O-Acetyl-L-serine hydrochloride | 66638-22-0 | Sigma | A6262 | 9.1 |
| 224 | Sodium hydroxide | 1310-73-2 | Mallinckrodt | 7708 | 42 |
| 225 | 5-Nitro-m-xylene-α,α′-diol | 71176-55-1 | Sigma | 184799 | 1.5 |
| 226 | 1,7-Heptanediol | 629-30-1 | Sigma | H2201 | 40 |
| 227 | L-(+)-Tartaric acid | 87-69-4 | Sigma | T1807 | 52 |
| 228 | N,N′-Dimethyl-1,3-propanediamine | 111-33-1 | Sigma | 308110 | 9.1 |

| | | | | | |
|---|---|---|---|---|---|
| 229 | 3,3′-Thiodipropionic acid | 111-17-1 | Sigma | T30201 | 2.9 |
| 230 | 1,4-Dimethylpyridinium p-toluenesulfonate | 78105-28-9 | Sigma | 514888 | 9.1 |
| 231 | sodium nitrite | 7632-00-0 | Sigma | S2252 | 40 |
| 232 | Triethyl citrate | 77-93-0 | Sigma | 27500 | 4.9 |
| 233 | 2-Methyl-1-propanol | 78-83-1 | Sigma | 294829 | 6.4 |
| 234 | Potassium iodide | 7681-11-0 | Fluka | 60399 | 52 |
| 235 | DL-Dithiothreitol | 3483-12-3 | Sigma | D5545 | 40 |
| 236 | DL-tartaric acid | 133-37-9 | Sigma | T400 | 14 |
| 237 | sodium cholate, hydrate | 206986-87-0 | Sigma | 270911 | 33 |
| 238 | sodium metabisulfite | 7681-57-4 | Sigma | S9000 | 35 |
| 239 | Sodium taurodeoxycholate hydrate | 207737-97-1 | Sigma | T0875 | 7.4 |
| 240 | Sodium iodide | 7681-82-5 | Sigma | 217638 | 62 |
| 241 | 1,4-Pentanediol | 626-95-9 | Sigma | 194182 | 50 |
| 242 | 2-Amino-5-diethylaminopentane | 140-80-7 | Sigma | A48806 | 9.1 |
| 243 | 2,2-Dimethyl-1,3-propanediol | 126-30-7 | Sigma | 538256 | 40 |
| 244 | Ethanolamine | 141-43-5 | Sigma | E0135 | 44 |
| 245 | trans-aconitic acid | 4023-65-8 | Sigma | 01600 | 24 |
| 246 | (±)-1,2,4-Butanetriol | 3068-00-6 | Sigma | 19040 | 50 |
| 247 | Aluminum sulfate hydrate | 7784-31-8 | Sigma | 227617 | 44 |
| 248 | 3-Piperidino-1,2-propanediol | 4847-93-2 | Sigma | 218499 | 9.1 |
| 249 | 1,3-Benzenedimethanol | 626-18-6 | Sigma | 196533 | 9.1 |
| 250 | 8-Aminooctanoic acid | 1002-57-9 | Sigma | 855294 | 20 |
| 251 | Magnesium chloride hexahydrate | 7791-18-6 | Sigma | M2670 | 33 |
| 252 | Calcium chloride dihydrate | 10035-04-8 | Sigma | C5080 | 11 |
| 253 | malic acid | 6915-15-7 | Sigma | M8304 | 31 |
| 254 | L-Cysteine hydrochloride monohydrate | 7048-04-6 | Sigma | C7880 | 3.9 |
| 255 | Zinc sulfate monohydrate | 7446-19-7 | Sigma | 307491 | 20 |
| 256 | 1,2,6-Hexanetriol | 106-69-4 | Sigma | T66206 | 50 |
| 257 | 1,3-Cyclopentanediol, mixture of cis and trans | 59719-74-3 | Sigma | 192805 | 50 |
| 258 | Camphor-10-sulfonic acid (β) | 5872-08-2 | Sigma | 147923 | 33 |
| 259 | succinic acid | 110-15-6 | Sigma | S9512 | 6.3 |
| 260 | Butylated hydroxyanisole | 25013-16-5 | Sigma | B1253 | 0.06 |

**ATCC reagent 18:**
0.75 g Trypticase Soy Broth
10 g Sucrose
5 g Bovine Serum Albumin Fraction V
100 mL Distilled water
Filter-sterilize through a 0.2 µm filter.

**ATCC reagent 20:**
20 g Sucrose
10 g Bovine Serum Albumin Fraction V
100 mL Distilled water
Filter-sterilize through a 0.2 µm filter.

**Table S3. Growth mediums used in this study**

*When a recipe is listed, details of the individual subcomponents are listed at end of table. Media used as a solid for plating assays was always solidified with 1.5% w/v agar.*

| Short name | Name or recipe | Vendor | Product # |
|---|---|---|---|
| *Final culture media used* | | | |
| Agar | Bacto Agar | BD | 214010 |
| LB | Difco LB Broth, Lennox | BD | 240230 |
| YPD | Difco YPD Broth | BD | 242820 |
| Nutrient | Difco Nutrient Broth | BD | 234000 |
| MRS | Difco Lactobacilli MRS Broth | BD | 288130 |
| MRS+cys | MRS + 0.05% w/v Cysteine | | |
| TY | per liter: 6g Tryptone, 3g Yeast extract, 0.5g CaCl2 | | |
| 1/2X BNM | per liter: 0.172g CaSO4, 0.195g MES,<br>2.5mL NOD major, 2.5mL NOD minor 1, 2.5mL NOD minor 2, 2.5 mL Fe-EDTA<br>pH to 6.5 with 2N KOH | | |
| *Subcomponents referenced in 1/2X BNM* | | | |
| NOD major | per liter: 24.4g MgSO4, 13.6g KH2PO4 | | |
| NOD minor 1 | per liter: 0.92g ZnSO4, 0.62g H3BO3, 1.69g MnSO4 | | |
| NOD minor 2 | per liter: 0.05g Na2MoO4, 0.0032g CuSO4, 0.005 CoCl2 | | |
| Fe-EDTA | per liter: 3.73g Na2EDTA, 2.78g FeSO4 | | |
| *Individual components referenced above* | | | |
| Tryptone | Bacto Tryptone | BD | 211705 |
| Yeast extract | Bacto Yeast Extract | BD | 212750 |
| CaCl2 | Calcium chloride dihydrate | Sigma | C5080 |
| Cysteine | L-Cysteine hydrochloride | Sigma | C7880 |
| CaSO4 | Calcium sulfate dihydrate | Sigma | C3771 |
| MES | MES hemisodium salt | Sigma | M8902 |
| MgSO4 | Magnesium sulfate heptahydrate | Sigma | 63138 |
| KH2PO4 | Potassium phosphate | Sigma | P5655 |
| ZnSO4 | Zinc sulfate heptahydrate | Sigma | Z1001 |
| H3BO3 | Boric acid | Sigma | B0394 |
| MnSO4 | Manganese (II) sulfate | Sigma | M7899 |
| Na2MoO4 | Sodium molybdate dihydrate | Sigma | M1651 |
| CuSO4 | Copper (II) sulfate | Sigma | C1297 |
| CoCl2 | Cobalt (II) chloride hexahydrate | Sigma | C2911 |
| Na2EDTA | EDTA disodium salt | Sigma | E6635 |
| FeSO4 | Iron (II) sulfate heptahydrate | Sigma | F8263 |
| KOH | Potassium hydroxide | Sigma | P5958 |

**Table S4. Strains and plasmids used in this study.**

| Strain | Source or Genotype | Note |
|---|---|---|
| *Escherichia coli* Nissle 1917 | Isolated from Mutaflor | Used in Fig. 2, S3, S4, S8, S9, |
| *Saccharomyces boulardii* | Isolated from Florastor | Used in Fig. 2, S3, S4 |
| *Ensifer meliloti* Rm1021 | ATCC 51124 | Used in Fig. 2, 4, S3, S4, S14 |
| *Lactobacillus plantarum* NC8 | CCUG 61730 | Used in Fig. 2, S3, S4 |
| *Medicago truncatula* A17 | Noble Research Institute | Used in Fig. 4 |
| sMJ026 | *E. coli* Nissle 1917 + pAKlux2 | Used in all other figures referencing *E. coli* Nissle 1917 |
| pAKlux2 (plasmid strain) | Addgene #14080 | Source of plasmid for sMJ026 |
